# Reduced *LANCL1-AS1* in old human skeletal muscle diminishes mitochondrial activity, shortens mt-mRNA poly(A) tails, and suppresses myogenesis

**DOI:** 10.64898/2026.07.05.736613

**Authors:** Jen-Hao Yang, Elizabeth K. Izydore, Krystyna Mazan-Mamczarz, Dimitrios Tsitsipatis, Julie A. Mattison, Brigette Romero, Changyou Shi, Xiaoling Yang, Rachel Munk, Jennifer L. Martindale, Carlos Anerillas, Martin Salamini-Montemurri, Martina Rossi, Yulan Piao, Jinshui Fan, Ya-Chen Chen, Bernardo Abel Cedeno-Veloz, Roberto Ferrero, Marta Montes, Nicolás Martínez-Velilla, Tian-Huei Chu, Kotb Abdelmohsen, Chang-Yi Cui, Mona Batish, Supriyo De, Payel Sen, Luigi Ferrucci, Rafael de Cabo, Myriam Gorospe

## Abstract

Regeneration of skeletal muscle preserves muscle mass and function, which decline with age. Here, we sought to identify long noncoding (lnc)RNAs involved in skeletal muscle myogenesis and potentially relevant to muscle aging. Cross-sectional analysis of skeletal muscle transcriptomes from healthy 22-through 89-year-old individuals revealed lncRNA *LANCL1-AS1* among the top declining transcripts. Conversely, *LANCL1-AS1* increased robustly during skeletal myogenesis and promoted myogenic differentiation in culture. Affinity pulldown by ChIRP followed by mass spectrometry revealed that *LANCL1-AS1* associated with the mitochondrial protein LRPPRC, enhancing the formation of the chaperone complex LRPPRC-SLIRP, which maintains longer poly(A) tails of mitochondrial (mt-)mRNAs and stabilizes mt-mRNAs. Importantly, while myoblasts from old rhesus monkey muscle expressed lower levels of *LANCL1-AS1* and mt-mRNAs, and displayed lower mitochondrial activity than young monkey myoblasts, overexpressing *LANCL1-AS1* in old myoblasts restored mitochondrial activity and myogenesis. We propose that the age-associated reduction in *LANCL1-AS1* contributes to impaired mitochondrial function and reduced myogenic capacity in aging skeletal muscle.

## INTRODUCTION

With advancing age, the skeletal muscle experiences a gradual loss of function resulting from declining number, quality, and size of muscle fibers.^1,2^ In turn, this loss of muscle is associated with frailty, fractures, and increased risk of metabolic diseases.^1,2^ In adult skeletal muscle, muscle mass is maintained primarily through the balance between protein synthesis and protein degradation, while myogenesis contributes to muscle growth during development and supports muscle regeneration following injury through the activity of muscle stem cells (satellite cells).^3^ As skeletal muscle ages, a decline in the numbers and activation of satellite cells, as well as changes in the cellular environment, may contribute to impaired regenerative capacity and, together with other age-associated alterations, to the progressive loss of muscle mass, strength, and function (sarcopenia).^4–6^ Given that muscle function, stem cell activity, and myogenic capacity are compromised with aging, there is mounting interest in developing strategies to preserve myogenesis and prevent sarcopenia.

A carefully orchestrated transcriptional program regulates myogenesis during muscle development and regeneration. Transcription factors (TFs) MYOD (myoblast determination protein 1) and MYF5 (myogenic factor 5) control the early stages of myoblast differentiation, followed by MYOG (myogenin), MRF4 (myogenic regulatory factor 4), and MEF2 (myocyte-specific enhancer factor 2), that drive subsequent stages of myogenesis.^7^ Besides TFs, a number of post-transcriptional regulatory RNA-binding proteins (RBPs) have been implicated in myogenesis, including ELAVL1 (embryonic lethal abnormal vision-like 1, also known as HuR), AUF1 (AU-binding factor 1), KHSRP (K-homology splicing regulatory protein), and CUGBP1 (CUG triplet repeat RNA-binding protein 1).^8–11^ These and other myogenesis-regulatory RBPs regulate the splicing, stability, and translation of key myogenic mRNAs.^12,13^

The other major regulators of myogenesis are noncoding RNAs. Long noncoding RNAs (lncRNAs, typically longer than 200 nucleotides) can interact with RNA, DNA, and proteins to control epigenetic alterations, transcription, RNA turnover, and translation.^14^ In this manner, lncRNAs regulate essential cellular processes including proliferation, differentiation, and survival; in turn, they orchestrate global processes such as muscle homeostasis.^15^ Besides interacting with RBPs, several myogenic lncRNAs associate with microRNAs (miRNAs), ∼22-nucleotide noncoding RNAs that repress target mRNAs with which they share partial complementarity.^16,17^ For instance, *lincMD1* controls myogenesis by sequestering miR-135 and miR-133a,^16^ *lnc-mg* promotes myogenesis and prevents muscle atrophy by neutralizing miR-125b,^17^ and *lncMyoD* promotes myoblast differentiation by suppressing the translation function of IMP2,^18^ while *lnc-31* enhances ROCK1 translation during myogenesis by binding to *Rock1* mRNA and YB-1.^19^ Mouse myogenesis is further stimulated by a rise in the circular RNA *circSamd4*, which inactivates transcriptional repressors PURA and PURB to enable *Myh* transcription.^20^ Human myogenesis is fostered by lncRNA *OIP5-AS1*, which scaffolds HuR and *MEF2C* mRNA, thereby enhancing the production of the myogenic transcription factor MEF2C.^21^ In parallel, *OIP5-AS1* lowers miR-7 levels by target-directed microRNA degradation, thus derepressing production of the fusogenic protein MYMX (myomixer), which is crucial for myoblast fusion and myotube generation.^22^ During human myogenesis, nuclear *lncFAM* accumulates and promotes differentiation by recruiting HNRNPL to the *MYBPC2* promoter, in turn increasing MYBPC2 levels.^23^

Here, we systematically investigated lncRNAs directly implicated in skeletal muscle aging. We examined the transcriptomes of skeletal muscle biopsies from a cross-sectional study of healthy individuals 22 to 89 years old to identify lncRNAs changing in expression levels. We compared them with the transcriptomes changing across human myogenesis in a cell culture model. The top candidate emerging from this comparison was *LANCL1-AS1*, which increased with myogenesis and declined with muscle aging. Interestingly, silencing *LANCL1-AS1* attenuated myogenesis and globally reduced the levels of mRNAs transcribed from the mitochondrial genome (mt-mRNAs). Molecular characterization revealed that *LANCL1-AS1* associated with the RBP LRPPRC, which, together with SLIRP, regulates mt-mRNA levels by preserving the length of their poly(A) tails and prolonging their half-lives. Accordingly, overexpressing *LANCL1-AS1*, but not a mutant *LANCL1-AS1* unable to interact with LRPPRC, restored myogenic capacity of primary myoblasts from muscle of old monkeys, where *LANCL1-AS1* is conserved. Supported by findings that sarcopenic human muscle exhibits lower *LANCL1-AS1* levels than age-matched controls, and that old skeletal muscle mt-mRNAs have shorter poly(A) tails than those in young muscle, we propose that *LANCL1-AS1* preserves myogenic capacity across the life span by enhancing mitochondrial function.

## RESULTS

### Human *LANCL1-AS1* levels decline in aging skeletal muscle and increase during myogenesis

To begin to understand the mechanisms underlying the age-associated loss of muscle mass and function, we recently reported the transcriptomes of human skeletal muscle biopsies from the healthy GESTALT cohort by RNA sequencing (RNA-seq) analysis.^24^ Identification of key functional processes altered in this cross-sectional study of 22- to 89-year-old (y.o.) participants^24^ using ShinyGO^25^ and the hallmark MsigDB database^26^ revealed that the top gene cluster reduced with age is associated with the category ‘myogenesis’ (Figure 1A), in keeping with evidence that muscle regeneration and myogenic capacity decline during aging.^27^ Heatmap visualization of changing lncRNAs from the GESTALT cohort (92 participants, STAR Methods), including the most significantly (adjusted p-value <0.05) declining and increasing lncRNAs with age, is shown (Figure 1B; Dataset 1). We then compared these transcriptomes with those of human cultured myoblasts AB1167 and AB678 undergoing myogenesis to differentiate into myotubes.^21,28^ RNA-seq analysis at the start of myogenesis (0 h), when myoblasts began to fuse (24 h), myotubes emerged (48 h), and myogenesis was fully established (72 h), revealed transcripts differentially abundant at different points^21,28^ (Figure 1C) [GSE202793^22^, GSE215343^23^, and GSE261476 (token oterkuicznajdij)]. Two lncRNAs were found to decline in human muscle aging and increase during myogenesis (Figure 1B-D; Datasets 1 and 2); given that during myogenesis *LINC00667* only increased modestly (Figure S1A) but *LANCL1-AS1* (*ENSG00000234281*, *AC007970.1*) increased dramatically, we focused on *LANCL1-AS1*, a lncRNA lacking coding potential (not shown).

**Figure 1.**
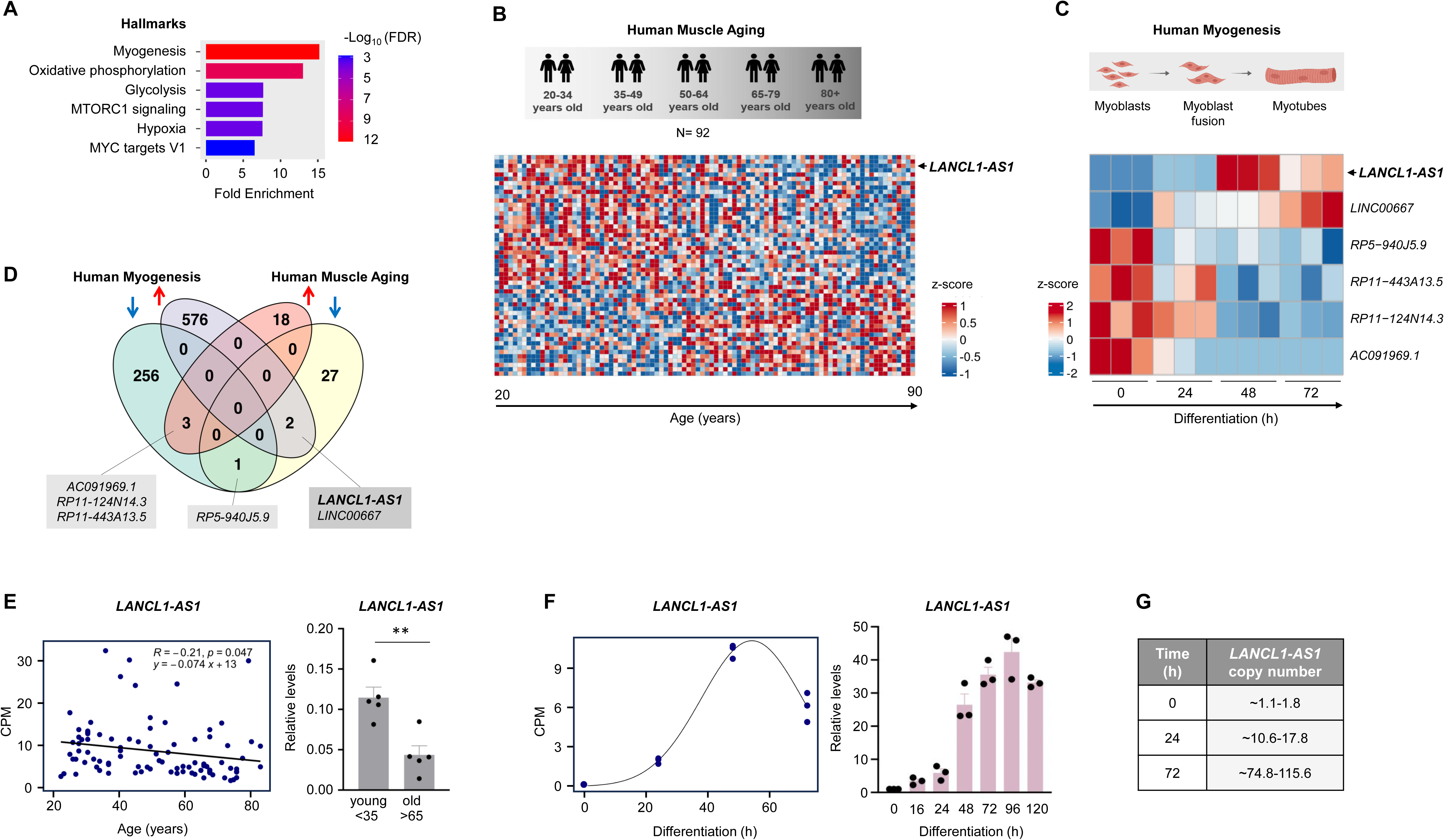
*LANCL1-AS1* levels decline with human muscle aging and increase with human myogenesis. **(A)** GSEA of pathways from the hallmark MSigDB database performed for the transcriptomes changing with age in human skeletal muscle (GESTALT study); myogenesis and oxidative phosphorylation are highlighted as those functional hallmarks declining most significantly. **(B)** Heatmap representation of lncRNAs (including antisense RNAs) differentially abundant with age in the GESTALT transcriptome dataset (92 participants total), at adjusted p-value <0.05, 51 RNAs (B); see also Dataset 1. **(C)** Following RNA-seq analysis of the transcriptomes of KM155 human myoblasts undergoing myogenesis (GSE261476, reviewer token oterkuicznajdij), those lncRNAs from (B) changing during myogenesis were identified and listed; total RNA was studied at the times of differentiation shown, 0 to 72 h (adjusted p-value <0.01) (C); see also Dataset 2. Top candidate lncRNA *LANCL1-AS1* is indicated; other lncRNAs shared between both datasets (human muscle aging and human myogenesis) are shown. **(D)** Venn diagram representation of the overlaps among lncRNAs significantly associated with human muscle aging and the transcriptomes from KM155 human myoblasts undergoing myogenesis (GSE261476, reviewer token oterkuicznajdij). Analysis included 51 RNAs from GESTALT (adjusted p-value <0.05; see also Dataset 1) and 838 RNAs from the myogenesis time course (adjusted p-value<0.01; see also Dataset 2). Among the 6 shared lncRNAs, *LANCL1-AS1* declined with muscle aging and increased during myogenesis; other 5 shared lncRNAs are indicated. **(E)** *Left*, expression levels of *LANCL1-AS1* per individual in the GESTALT cohort by linear fit plot representation; log_2_ (CPM) values from the RNA-seq datasets from muscle biopsies;^24^ *right*, validation of *LANCL1-AS1* levels in RNA from fresh muscle biopsies in the same study^24^ by RT-qPCR analysis, normalized to *GAPDH* mRNA levels. **(F)** *Left*, expression levels of *LANCL1-AS1* per individual myoblast sample by non-linear fit plot representation; log_2_ (CPM) values from the RNA-seq datasets from myoblasts undergoing differentiation (GSE164471; GSE261476, reviewer token oterkuicznajdij]);^24^ *right*, validation of *LANCL1-AS1* levels by RT-qPCR analysis in freshly prepared RNA from myoblasts undergoing myogenesis (collected at the times shown), normalized to *GAPDH* mRNA levels. **(G)** Quantification of copy numbers of *LANCL1-AS1* in AB678 myoblasts (STAR Methods) during myogenesis, shown as number of RNA molecules per cell (time 0) or per nucleus equivalent (times 24 h and 48 h). Data in (E, F) are the means ± SEM from three or more biological replicates. Significance (*, p<0.05; **, p<0.01; ***, p<0.001) was established using Student’s *t*-test.

The levels of *LANCL1-AS1* declined gradually and significantly with advancing age (Figure 1E, *left*), and reverse transcription followed by quantitative PCR analysis (RT-qPCR) of RNA freshly extracted from muscle biopsies of young (<35 y.o.) and old (>65 y.o.) individuals (n=5) from the GESTALT cohort further supported the conclusion that *LANCL1-AS1* levels significantly decreased in the older group (Figure 1E, *right*). Conversely, the levels of *LANCL1-AS1* across myogenesis, visualized both from the RNA-seq analysis in Figure 1C (Figure 1F, *left*) and by RT-qPCR analysis of fresh RNA isolated from cultured myoblasts (bar graph, Figure 1F, *right*), revealed an early rise in the levels of *LANCL1-AS1* at the start of myogenesis (24 h), which further increased during fusion into myotubes (48-72 h) and in fully established differentiated cultures (96 h). This pattern of expression in differentiating myoblasts suggested that *LANCL1-AS1* may play an important role in late stages of myogenesis. In fact, a limited survey of other cultured cells suggested that *LANCL1-AS1* is more abundant in neuronal and skeletal muscle cells, although it was detected in other organs such as lung and thyroid (Genecards.org); *MYL1* mRNA, a control myogenic transcript, was also assessed (Figure S1B). *LANCL1-AS1* did not appear to regulate *MYL1* transcription through direct binding to its promoter (not shown).

In myoblasts, the sequence of *LANCL1-AS1*, established using 5’ RACE (STAR Methods), varied slightly from the reported NCBI reference sequence (NR_110604.1), as it lacked the 9 terminal nucleotides and contained an additional 175 nucleotides inserted near the 5’ end (Figure S1C and S1D). Additionally, the number of *LANCL1-AS1* copies was calculated using droplet digital PCR analysis, by interpolating on a standard curve of known nucleic acid concentrations, and by comparing relative Ct values (STAR Methods)^29^; as shown, *LANCL1-AS1* transcripts ranged from ∼1-2 copies per cell in proliferating myoblasts to ∼85–115 copies per nucleus equivalent in differentiated myotubes (Figure 1G, Figure S1E). In summary, the lncRNA showing the greatest decline with age in human skeletal muscle, *LANCL1-AS1*, was found to increase robustly during human myogenesis, suggesting that it might be part of the program of muscle preservation in young persons that becomes gradually less efficient with aging.

### Silencing *LANCL1-AS1* suppresses myogenesis

To investigate the function of *LANCL1-AS1*, we first used the human myoblast line AB678 to examine the impact of reducing *LANCL1-AS1* levels on myogenic differentiation. Myoblasts proliferating in growth media (GM) were cultured to high density and switched to differentiation media (DM, supplemented with 2% horse serum) to trigger myogenesis, and gradually fuse into myotubes over the following 48 to 72 h; this process was visualized by immunofluorescence detection of the myogenic differentiation marker MYH (Figure 2A). We reduced *LANCL1-AS1* levels in AB678 myoblasts by transfecting siRNAs directed at *LANCL1-AS1*; Ctrl siRNA was used in control transfections (Figure 2B). Importantly, silencing *LANCL1-AS1* 24 h before initiating myogenesis potently reduced and delayed myogenic progression, as determined by MYH immunofluorescence analysis to track myotube formation, as thinner and less numerous myotubes appeared at later times after silencing *LANCL1-AS1* (Figure 2C). Quantifying myotubes further revealed that silencing *LANCL1-AS1* reduced the fusion index (the number of nuclei inside MYH-positive myotubes relative to the total number of nuclei) from ∼90% to ∼30% by 72 h into differentiation, and that the average number of nuclei per myotube declined from ∼35 to ∼5 (Figure 2D). Experiments using another LANCL1-AS1 siRNA (#2) and another human myoblast line, AB1167, showed similar results (Figure S2).

**Figure 2.**
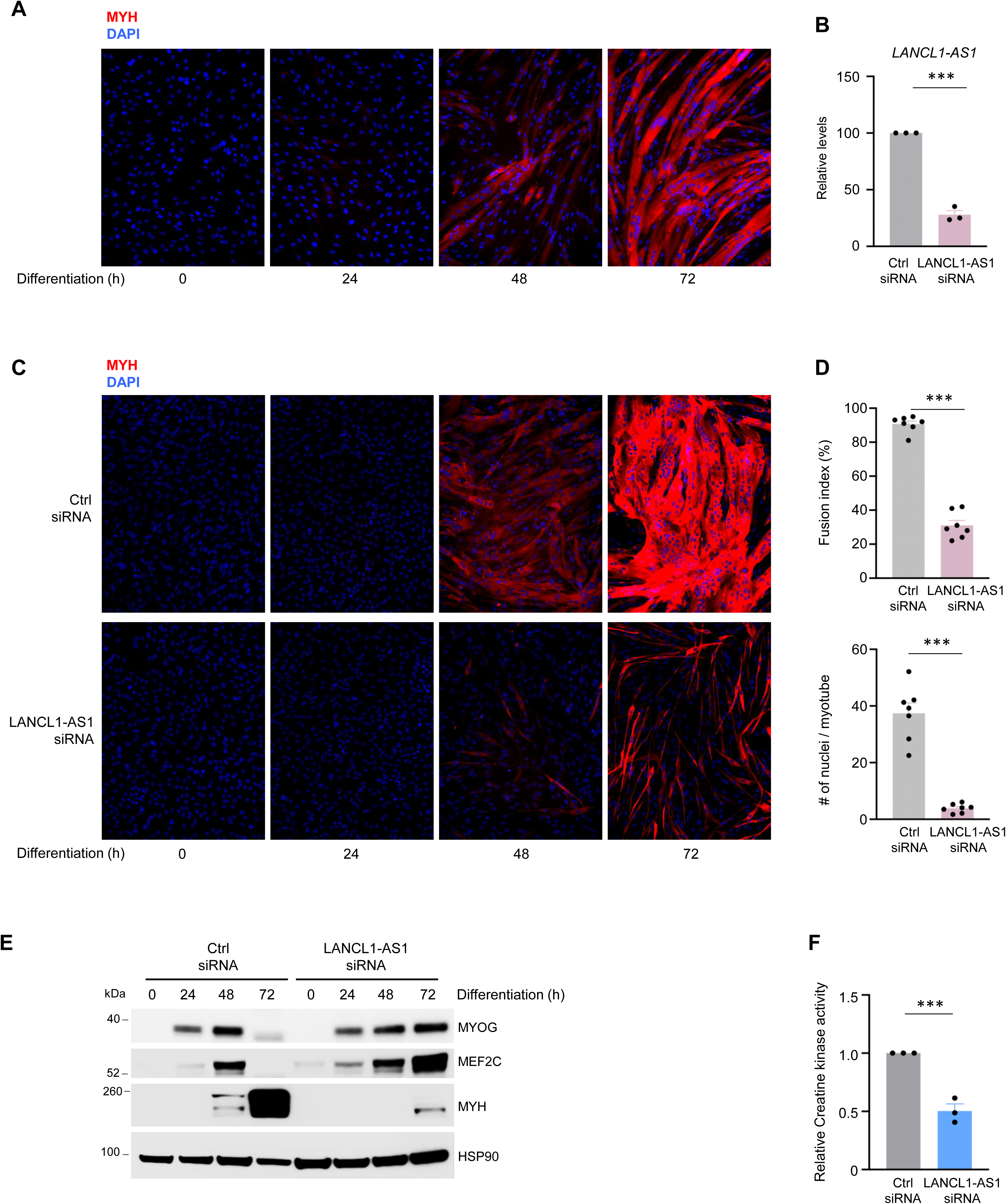
Silencing *LANCL1-AS1* reduces human myogenesis. **(A)** Immunofluorescence analysis of MYH signals (red) to monitor the progression of human AB678 myoblasts from a proliferative state (0 h), when MYH is undetectable, to a state of full differentiation into myotubes (72 h), when MYH fluorescence is robust. DAPI (blue) was used to visualize nuclei. **(B)** Proliferating AB678 myoblasts were transfected with Ctrl siRNA or LANCL1-AS1-directed siRNA #1; 48 h later, the levels of remaining *LANCL1-AS1* were measured by RT-qPCR analysis. *GAPDH* mRNA levels were also quantified for normalization. **(C, D)** AB678 myoblasts were transfected with Ctrl siRNA or LANCL1-AS1 siRNA #1; 24 h later, they were placed in differentiation medium for indicated times, whereupon differentiation was monitored by assessing MYH levels (red) by immunofluorescence (C). The fusion index and number of nuclei (stained using DAPI) per myotube after differentiation for 72 h were quantified (D); five separate fields were evaluated per experiment. **(E)** The levels of myogenic proteins MYOG, MEF2C, MYH, and loading control HSP90 were assessed by western blot analysis of lysates from AB678 myoblasts that were transfected and processed as explained in (C); lysates were collected at the times shown after the induction of differentiation. **(F)** Cells were processed as described in (C), and the levels of creatine kinase (CK) activity at 72 h of differentiation were measured enzymatically (STAR Methods). Data in (B,D,F) represent the means ± SEM from three or more biological replicates. Significance (*, p<0.05; **, p<0.01; ***, p<0.001) was established using Student’s *t*-test.

In keeping with these phenotypes and the relatively late increase in *LANCL1-AS1* levels during myogenesis (Figure 1F), silencing *LANCL1-AS1* robustly reduced the levels of the late differentiation marker MYH, but only modestly affected the earlier differentiation markers MYOG and MEF2C (Figure 2E), which remained elevated at 72 h, with ongoing myotube formation. Finally, silencing *LANCL1-AS1* reduced creatine kinase activity, another indicator of myogenic differentiation, at 72 h into myogenesis (Figure 2F). Together, these results mirror the low levels of *LANCL1-AS1* early in myogenesis (Figure 1F) and support the notion that *LANCL1-AS1* is important later in myogenesis, when it is more abundant.

### Silencing *LANCL1-AS1* reduces mitochondrial activity

To study the molecular mechanisms through which *LANCL1-AS1* regulates myogenesis, we first investigated how silencing *LANCL1-AS1* in AB678 cells influenced myogenic transcriptomic programs. We initiated myogenic differentiation 24 h after transfecting Ctrl or LANCL1-AS1 siRNAs, then 72 h later we performed RNA-seq analysis [GSE215343^23^ and GSE261477 (token yzqxkuykfvopbqx)]. Gene Set Enrichment Analysis (GSEA) of GO Biological Process dataset terms revealed that mRNAs related to muscle contraction, development, and ion transport markedly decreased after silencing *LANCL1-AS1* (Figure 3A,B). Given that myogenesis and muscle contraction are intimately linked to mitochondrial activity and ion channel function,^30–33^ and considering that the second functional gene cluster declining with human muscle aging was oxidative phosphorylation (Figure 1A), we set out to investigate if *LANCL1-AS1* influences mitochondrial function during human myogenesis.

**Figure 3.**
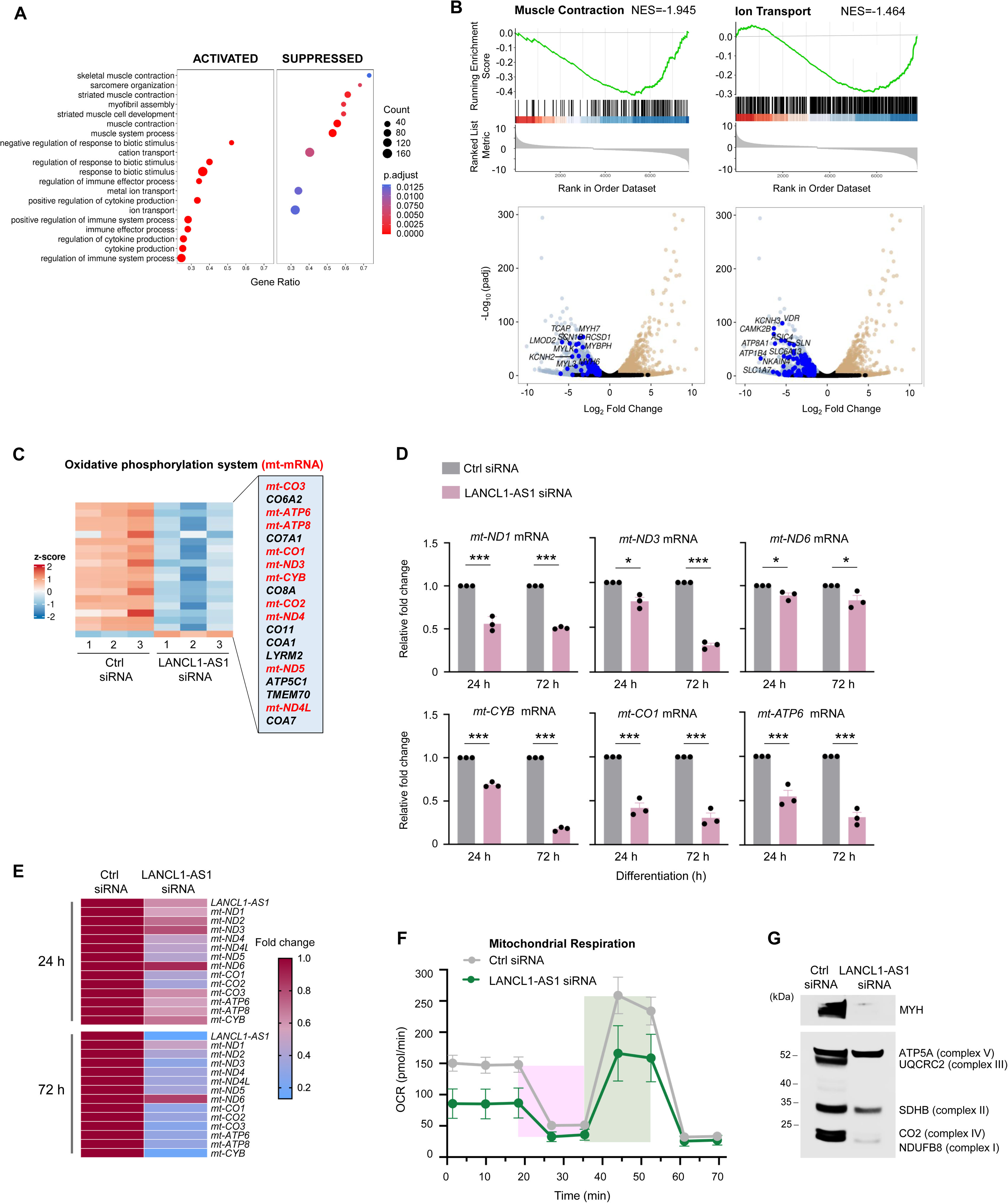
Loss of *LANCL1-AS1* attenuates mitochondrial function, reduces abundance of mt-mRNAs. **(A)** AB678 myoblasts were transfected with Ctrl siRNA or LANCL1-AS1 siRNA; 24 h later, they were placed in differentiation medium for 72 h, whereupon RNA was collected for RNA-seq analysis [GSE261477 (token yzqxkuykfvopbqx)]. Top 20 significantly activated and suppressed functional networks of the Gene Ontology Biological Process dataset obtained by GSEA of the mRNAs differentially expressed after silencing *LANCL1-AS1* compared to control populations. ‘Gene Ratio’ represents the proportion of those mRNAs significantly changing from the total number of mRNAs implicated in a given functional network. **(B)** Analysis of the transcriptomes in AB678 cells processed as in (A); GSEA associations (*top*), and volcano plots (*bottom*) depict genes encoding mRNAs related to muscle contraction and ion channel function that changed significantly in cells after silencing *LANCL1-AS1*. **(C)** Heatmap representation of changes in the transcriptome associated with ‘Oxidative phosphorylation system’ after silencing *LANCL1-AS1*, including mRNAs encoded by the nuclear genome (black) and mRNAs encoded by the mitochondrial genome (mt-mRNAs, red); triplicate samples are shown [GSE261477 (token yzqxkuykfvopbqx)]. See Figure S3 for complete heatmap representation of other systems changing in this analysis. **(D)** AB678 myoblasts were transfected with Ctrl siRNA or LANCL1-AS1 siRNA; 24 h later, they were placed in differentiation medium and collected at 24 and 72 h after inducing differentiation. The levels of mitochondrial mRNAs *mt-ND1*, *mt-ND3*, *mt-ND6*, *mt-CYB*, *mt-CO1*, and *mt-ATP6* mRNAs were measured by RT-qPCR analysis, normalized to *GAPDH* mRNA levels. **(E)** Heatmap representation of the levels of mt-mRNAs and *LANCL1-AS1* [GSE261477 (token yzqxkuykfvopbqx)] in AB678 cultures processed as described in (D). **(F)** Measurement of mitochondrial respiration of AB678 cultures at 24, 48, and 72 h of differentiation using a Seahorse XFe24 analyzer. Green shaded region, maximal respiration; pink shaded region, ATP production. **(G)** AB678 myoblasts were transfected with Ctrl or LANCL1-AS1 siRNAs; 24 h later, they were placed in differentiation medium for 72 h. The levels of MYH, as well as proteins in the oxidative phosphorylation complexes I (NDUFB8), II (SDHB), III (UQCRC2), IV (COX2), and V (ATP5A), and loading control HSP90 were assessed by western blot analysis. Data in (D, F) are the means +SD of three or more independent experiments. Significance (*, p<0.05; **, p<0.01; ***, p<0.001) was established using Student’s *t*-test.

We first studied if silencing *LANCL1-AS1* influenced mitochondrial gene expression and pathways utilizing MitoCarta 3.0 datasets.^34^ Interestingly, the levels of many mRNAs transcribed from the mitochondrial genome (mt-mRNAs) were affected by silencing *LANCL1-AS1*—with some increasing and some decreasing (Figure S3A). Intriguingly, however, those mRNAs encoding proteins with roles in oxidative phosphorylation were almost all reduced after silencing *LANCL1-AS1*, and many of them were mt-mRNAs (Figure 3C, Figure S3B). In agreement with the RNA-seq results, silencing *LANCL1-AS1* lowered the levels of mt-mRNAs at 24 and 72 h of differentiation, as quantified by RT-qPCR analysis.

Among them, the reduction in levels of *mt-ND1*, *mt-ND2*, *mt-ND3*, *mt-CO3*, *mt-ATP6*, *mt-ATP8*, and *mt-CYB* mRNAs was modest at 24 h of differentiation and was more pronounced at 72 h of differentiation. Notably, *mt-ND6* mRNA only changed slightly at both 24 and 72 h of differentiation (Figure 3D,E).

Considering that mitochondrial activity increases during myogenesis,^35^ we examined if mitochondrial function (as assessed by evaluating oxygen consumption rate, OCR) changed during human myogenesis. As shown in Figure S3C, mitochondrial activity increased during myogenesis, with maximal respiration measured at 72 h of differentiation (Figure S3C, green shade), while ATP production capacities increased both at 24 and 72 h of differentiation (Figure S3C, pink shade). Moreover, silencing *LANCL1-AS1* reduced all parameters of mitochondrial activity (including basal level, spare capacity, maximal respiration, and ATP production) at 72 h of differentiation (Figure 3F, but not mt-DNA levels, not shown). We extended these findings to elucidate if protein components of oxidative phosphorylation complexes changed after silencing *LANCL1-AS1*; as shown, silencing *LANCL1-AS1* strongly reduced the levels of several such proteins, including complex I protein NDUFB8, complex II protein SDHB (encoded by the nuclear genome), complex III protein UQCRC2, and complex IV protein COX II (Figure 3G). These results connect *LANCL1-AS1* levels to mitochondrial activity during myogenesis and with aging.

### *LANCL1-AS1* binds mitochondrial RBPs LRPPRC and SLIRP

LncRNA function is often associated with the RBPs they bind.^14,18,36^ To investigate the RBPs interacting with *LANCL1-AS1*, we performed ChIRP-MS (comprehensive identification of RNA-binding proteins by mass spectrometry). Differentiated myoblasts were crosslinked using 3% formaldehyde, lysed, and homogenized by sonication. A set of 14 biotinylated antisense oligonucleotides (ASOs) designed to bind to *LANCL1-AS1* were then added to the lysate, enabling the specific capture of *LANCL1-AS1*-interacting complexes using magnetic streptavidin-coated beads (Figure 4A). The pulldown efficiency was evaluated (Figure S4A) and the interacting proteins identified by mass spectrometry analysis (MS, Table S1). LacZ ASOs were used in control pulldown reactions. We identified a total of 42 proteins selectively interacting with *LANCL1-AS1*, many of them mitochondrial proteins (Figure 4B, Table S1), suggesting that *LANCL1-AS1* is at least partially present in mitochondria. Notably, the top candidate protein was the mitochondrial RBP LRPPRC (Figure 4B); LRPPRC is encoded by a gene that is mutated in Leigh syndrome French-Canadian type, associated with mitochondrial defects^37–39^ LRPPRC is implicated in the post-transcriptional regulation of the expression of proteins encoded by the mitochondrial genome, and is implicated in energy metabolism and mitochondrial function, prompting us to investigate LRPPRC further.

**Figure 4.**
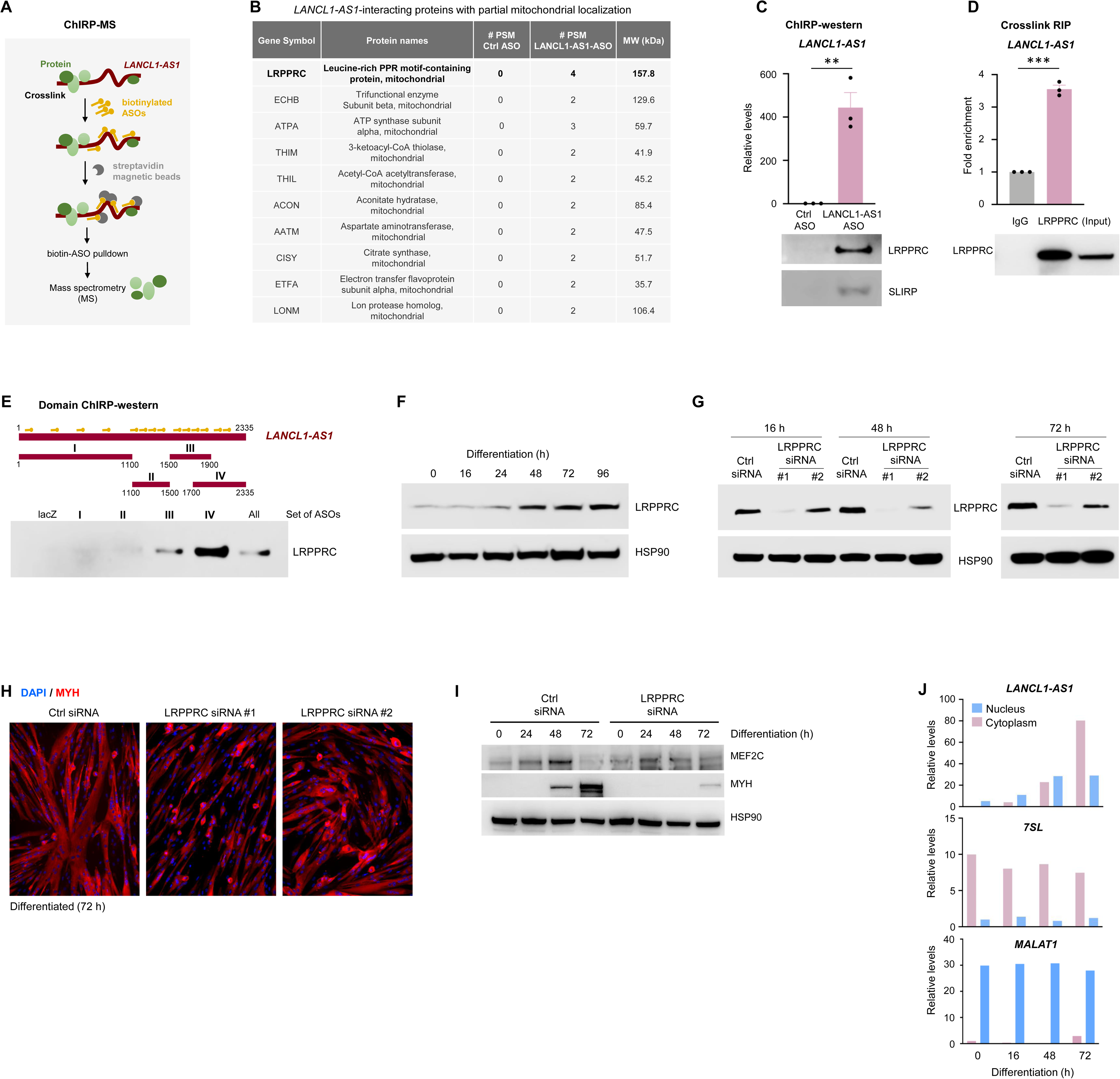
*LANCL1-AS1* binds mitochondrial RBP LRPPRC. **(A)** Schematic of the ChIRP-MS workflow. Briefly, AB678 cultures undergoing differentiation for 72 h were crosslinked with 3% formaldehyde for 30 min and sonicated. After incubating the sonicated lysates with biotinylated antisense oligomers (ASOs) directed at *LANCL1-AS1* or at control *LacZ* mRNA, complexes were pulled down using streptavidin-coated magnetic beads. Interacting proteins were eluted for analysis by mass spectrometry. **(B)** Top 10 *LANCL1-AS1*-interacting proteins with at least partial mitochondrial localization, identified from differentiated AB678 cells by ChIRP-MS analysis. **(C)** After performing ChIRP analysis as described in (A), RT-qPCR analysis was used to measure the enrichment in *LANCL1-AS1* in the pulldown materials using biotinylated ASOs (relative to control LacZ biotin ASO pulldown) (*top*), and western blot analysis was performed to detect *LANCL1-AS1*-interacting protein LRPPRC, and the LRPPRC partner protein SLIRP (*bottom*). **(D)** Analysis of the interaction between *LANCL1-AS1* and LRPPRC by crosslinking RNP immunoprecipitation (IP) assays using anti-LRPPRC or control IgG antibodies. The enrichment of *LANCL1-AS1* in the IP was determined by RT-qPCR analysis after normalizing to *GAPDH* mRNA levels (*top*), and the presence of LRPPRC in the IP was assessed by western blot analysis; ‘*Input*’, lysate aliquot *(bottom)*. **(E)** ChIRP-western experiment performed as described in (C), but incubating the sonicated RNA (∼500-nt fragments) with different pools of ASOs. ASOs 1-4 (I), 5-8 (II), 9-12 (III), 11-14 (IV), and all ASOs (1-14) with the same final concentration (nM) of ASOs for each segment. The levels of LRPPRC after performing ChIRP with different pools of ASOs directed at different segments (I through IV) of *LANCL1-AS1* were assessed by western blot analysis. **(F)** Levels of LRPPRC and loading control HSP90, as assessed by western blot analysis in AB678 cultures undergoing differentiation for the times indicated. **(G-I)** AB678 myoblasts were transfected with Ctrl siRNA or LRPPRC-directed siRNAs #1 or #2; 24 h later, they were placed in differentiation medium for an additional 16, 48, or 72 h; the levels of LRPPRC and loading control HSP90 were assessed by western blot analysis (G). Myogenic progression was monitored by assessing MYH immunofluorescent signals by 72 h into differentiation (H). The levels of MEF2C, MYH, and loading control HSP90 were assessed by western blot analysis at the times indicated (I). **(J)** Following fractionation of nuclear and cytoplasmic lysates from AB678 cultures that were proliferating or undergoing myogenic differentiation for 16, 48, or 72 h, the levels of *LANCL1-AS1* (and cytoplasmic and nuclear control lncRNAs, *7SL* and *MALAT1*, respectively) were measured by RT-qPCR analysis. Data in (C, D) represent the means ±SEM of three or more biological replicates. Significance (**, p<0.01; ***, p<0.001) was established using Student’s *t*-test.

We validated the interaction between *LANCL1-AS1* and LRPPRC by performing ChIRP followed by western blot analysis. Pulldown with *LANCL1-AS1* and Ctrl ASOs was used to confirm the enrichment of *LANCL1-AS1* by RT-qPCR analysis (Figure 4C, *top*) and LRPPRC by western blot analysis (Figure 4C, *bottom*). Interestingly, LRPPRC functions together with another mitochondrial RBP, SLIRP.^40^ SLIRP is a small protein (∼100 amino acids long) and is thus difficult to capture in large-scale ChIRP-MS; however, we were able to validate its interaction with *LANCL1-AS1* by ChIRP-western blot analysis (Figure 4C, *bottom*). These validation efforts were important, as some presumed *LANCL1-AS1*-interacting proteins from ChIRP-MS were not detected in the pulldown material (not shown).

For further validation, we used an orthogonal approach consisting of crosslinking followed by ribonucleoprotein (RNP) immunoprecipitation (IP), to confirm the interactions between LRPPRC and *LANCL1-AS1*. Briefly, at 72 h into differentiation, AB678 cultures were crosslinked and RNP IP (RIP) analysis was performed using an antibody recognizing LRPPRC (or IgG in control IP reactions); the bound RNAs were then quantified by RT-qPCR analysis. As shown, *LANCL1-AS1* was highly enriched in LRPPRC IP samples, but not in control IgG IP samples (Figure 4D, *top*). The successful IP of LRPPRC was confirmed by western blot analysis of the IP samples (Figure 4D, *bottom*).

To identify the region(s) of interaction between LRPPRC and *LANCL1-AS1*, we carried out domain ChIRP-western analysis. Similar to domain ChIRP for identifying RNA-chromatin interactions, the ASOs were split into 4 groups, with each group corresponding to different regions of *LANCL1-AS1*. After sonication to fragment the RNA into ∼500-nt transcript segments, the RNA was incubated with each set of region-specific ASOs; following pulldown, western blot analysis was used to identify the region of *LANCL1-AS1* associating with LRPPRC. As shown, pulldown with ASOs 11-14 (segment IV) showed the strongest binding between *LANCL1-AS1* and LRPPRC (Figure 4E); ASOs 9-12 (segment III) showed substantially less binding. Notably, the combined pool of 14 (‘All’) ASOs pulled down LRPPRC with lower strength; this is because the final concentration of total ASOs was kept constant, and therefore ASOs 11-14 in the mix were relatively less abundant. The pulldown efficiency of these domain-specific ASOs was very high (∼400-fold enrichment of *LANCL1-AS1* in *LANCL1-AS1* pulldown experiment), similar to the enrichment seen for *LANCL1-AS1* without RNA fragmentation (Figure S4A and S4B). These results strongly support the notion that the 3’ distal end of *LANCL1-AS1* is the primary region of interaction with LRPPRC.

We then sought to elucidate the function of the *LANCL1-AS1*-LRPPRC lncRNP complex in myogenesis and muscle aging. Analysis of LRPPRC protein levels in AB678 cultures by western blot analysis revealed that LRPPRC levels increased during myogenic differentiation (Figure 4F). Interestingly, silencing LRPPRC by transfection of two different siRNAs directed at *LRPPRC* mRNA between 16 and 72 h in differentiation medium (Figure 4G) diminished myotube formation, as determined by evaluating MYH immunofluorescence (Figure 4H), but did not affect metabolic pathways or cell survival (not shown). Analysis of protein markers of myogenesis revealed that silencing LRPPRC delayed slightly the rise in production of the early myogenic markers MEF2C (Figure 4I) and MYMX (not shown), but potently reduced the appearance of the late differentiation marker MYH by 72 h of differentiation (Figure 4I).

We further investigated this complex by studying its localization in the cell. First, we prepared nuclear and cytoplasmic fractions (STAR Methods) and measured the abundance of *LANCL1-AS1* in each compartment by RT-qPCR analysis; control lncRNAs that preferentially localized in the cytoplasm (*7SL*) and nucleus (*MALAT1*) were also monitored to ensure a proper subcellular fractionation (Figure 4J). Despite its low abundance, *LANCL1-AS1* was seen in the nucleus of proliferating myoblasts; however, during MYH synthesis in differentiating myotubes *LANCL1-AS1* is robustly mobilized to the cytoplasm (Figure 4J). Single-molecule fluorescence *in situ* hybridization analysis (smFISH, STAR Methods) further confirmed that *LANCL1-AS1* was mainly nuclear in proliferating myoblasts and predominantly cytoplasmic (and markedly more abundant) in myotubes (Figure 5A). The quantification of smFISH signals confirmed that *LANCL1-AS1* is primarily localized in the nucleus of proliferating myoblasts; upon differentiation into myotubes, its expression increases dramatically, accompanied by a shift to a predominantly cytoplasmic distribution (Figure 5B and Figure S4C). In agreement with earlier reports,^40^ immunofluorescence data revealed that LRPPRC largely colocalized with the mitochondrial protein VDAC, indicating that LRPPRC is mainly found in mitochondria (Figure 5C). Further, smFISH signals for *LANCL1-AS1* and immunofluorescence signals to detect mitochondrial markers MitoTracker, TOM20, and LRPPRC revealed that LRPPRC interacts with *LANCL1-AS1* at least partly in mitochondria (Figure 5D and Figure S4D and E). The overlapping signals were further quantified (Figure 5E) and revealed that approximately 20% of *LANCL1-AS1* colocalized with mitochondrial markers. Signal overlaps were visualized with yellow circles (Figure S4E); as a control, we included a known nuclear lncRNA, *LncFAM* (Figure S4E).^23^

**Figure 5.**
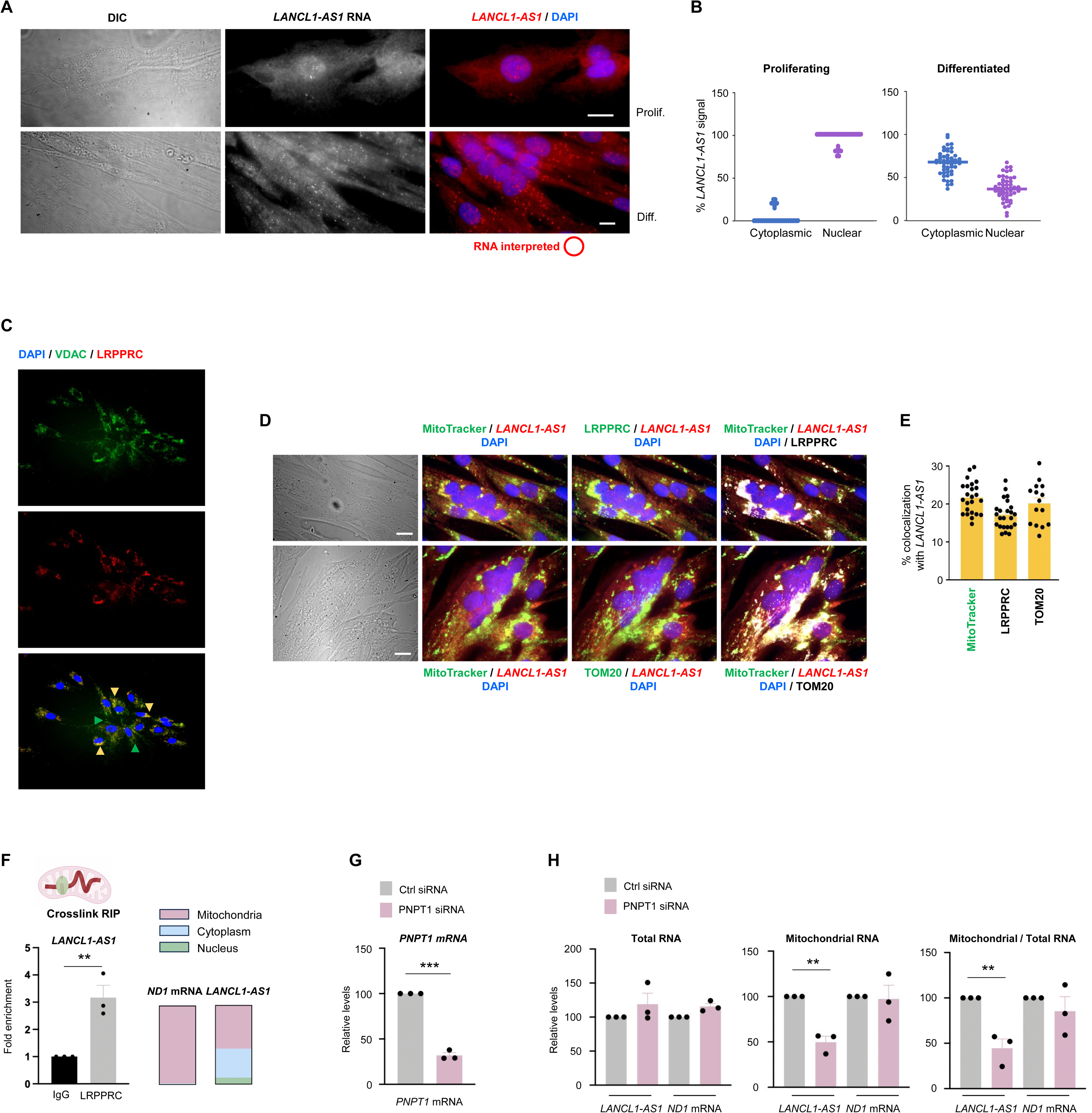
Partial localization of *LANCL1-AS1* in mitochondria and import role or PNPT1. **(A)** Representative smFISH performed in AB678 cultures using a probe set for *LANCL1-AS1* RNA labeled with Texas red; *left*, differential interference contrast (DIC) micrographs; *center*, raw merged Z stacks of RNA signal (gray); *right*, RNA signal pseudocolored red, merged with blue pseudocolored DAPI stain and overlaid with red circles to highlight the RNA spots identified using MATLAB. Scale bar, 5 µm. **(B)** Quantification of the relative levels of nuclear and cytoplasmic *LANCL1-AS1* smFISH signals in AB678 cultures that were proliferating or had differentiated for 72 h; ∼50 myotubes were evaluated in three biological repeats. **(C)** Immunofluorescence analysis of the colocalization of signals for VDAC (green) and LRPPRC (red) in differentiated (72 h) AB678 cells. Blue, nuclei stained with DAPI; yellow arrows, cells co-stained for VDAC and LRPPRC; green arrows, VDAC-only positive cells. **(D)** Representative smFISH analysis of *LANCL1-AS1* colocalizing with MitoTracker, LRPPRC, TOM20. *Left*, DIC micrographs. *Left center*, RNA signals using probes for *LANCL1-AS1* RNA (red), merged with MitoTracker (green) and DAPI (blue) signals. *Right center*, RNA signals using probes for *LANCL1-AS1* RNA (red), merged with LRPPRC (*top*) or TOM20 (*bottom*) signals using antibodies against LRPPRC or TOM20 (green), and DAPI (blue); light orange spots denote where lncRNA signals colocalize with LRPPRC or TOM20. *Right,* RNA signals using probes for *LANCL1-AS1* RNA (red), merged with LRPPRC (*top*) or TOM20 (*bottom*) signals using antibodies against LRPPRC or TOM20 (white), and DAPI (blue), further merged with MitoTracker signals (green) and DAPI (blue); light orange spots denote places where lncRNA signals colocalize with both MitoTracker and LRPPRC or MitoTracker and TOM20 as identified using MATLAB. Scale bar, 5 µm. **(E)** Quantification of smFISH *LANCL1-AS1* signals colocalizing with MitoTracker, LRPPRC, or TOM20. **(F)** After differentiation for 72 h, AB678 cultures were crosslinked with 0.1% formaldehyde and mitochondria were isolated for and RIP analysis (STAR Methods) using anti-LRPPRC and control IgG antibodies (*left*). The enrichment of *LANCL1-AS1* associated with LRPPRC was measured by RT-qPCR analysis. In the same preparations (AB678 cultures differentiated for 72 h, then crosslinked), the levels of *mt-ND1* mRNA and *LANCL1-AS1* were measured by RT-qPCR analysis in mitochondrial, cytoplasmic, and nuclear fractions. **(G,H)** AB678 myoblasts were transfected with Ctrl siRNA or siRNAs directed at *PNPT1* mRNA; 24 h later, they were placed in differentiation conditions, and after an additional 72 h the levels of *PNPT1* mRNA were assessed by RT-qPCR analysis (G), and the levels of mitochondrial *LANCL1-AS1* RNA, Total *LANCL1-AS1* RNA, and Mitochondrial RNA relative to Total *LANCL1-AS1* RNA (H), were assessed by RT-qPCR analysis. Data in (F-H) represent the means ±SEM of three or more biological replicates. Significance (**, p<0.01; ***, p<0.001) was established using Student’s *t*-test.

To complement these findings, we studied if LRPPRC associated with *LANCL1-AS1* in mitochondria of myotubes. After crosslinking of differentiated cultures with formaldehyde, isolating mitochondria, and testing by the interaction of LRPPRC with *LANCL1-AS1* by RIP analysis, we found a significant enrichment in *LANCL1-AS1* binding to LRPPRC (Figure 5F, *left*). Quantification of *LANCL1-AS1* in different cell compartments indicated that approximately one-half of *LANCL1-AS1* was found in mitochondria of differentiated myotubes, and the other half was primarily cytoplasmic with a small proportion of *LANCL1-AS1* in the nucleus (Figure 5F, *right*), in agreement with the smFISH data (Figure 5B). The translocation of lncRNAs from the cytoplasm to mitochondria is poorly understood, but PNPase (PNPT1) has been reported to assist with RNA import into mitochondria.^41,42^ We thus examined if PNPT1 regulated the import of *LANCL1-AS1* into mitochondria. After silencing PNPT1 by transfecting siRNAs directed at *PNPT1* mRNA in AB678 cells (Figure 5G), and inducing differentiation for 72 h, we isolated total RNA and mitochondrial RNA. RT-qPCR analysis to assess the levels of *LANCL1-AS1* and *mt-ND1* mRNA revealed that silencing *PNPT1* mRNA (Figure 5H) did not affect cell morphology (Figure S4F) or the steady-state levels of either *LANCL1-AS1* or *mt-ND1* mRNA (Figure 5H, *left graph*). However, silencing *PNPT1* mRNA reduced the levels of *LANCL1-AS1* in the mitochondria, while the levels of mitochondrial *mt-ND1* mRNA remained unaffected (Figure 5H), supporting the notion that PNPT1 serves, at least in part, to import *LANCL1-AS1* into mitochondria. Together, these results indicate that human *LANCL1-AS1* interacts with LRPPRC in mitochondria, supporting a possible function for this lncRNP complex in mitochondrial activity and human myogenesis.

### The *LANCL1-AS1*-LRPPRC lncRNP enhances polyadenylation and stabilization of mt-mRNAs

Although silencing *LANCL1-AS1* globally reduced mt-mRNA levels, this effect was independent of mtDNA biosynthesis, as mtDNA levels remained unchanged (a representative segment, *mt-ND1* DNA, shown in Figure S5A), suggesting that the regulation may occur independently of transcription. LRPPRC is a multifunctional protein involved in maintaining mt-RNA poly(A) tail length to improve mt-mRNA stability and translation; the resulting enhancement in production of mitochondrial proteins promotes both mitochondrial function and energy production.^39,40^ To investigate if the *LANCL1-AS1*-LRPPRC complex contributed to the stability of mt-mRNAs, we measured their half-lives. Briefly, *LANCL1-AS1* was silenced in AB678 cells by transfecting siRNAs at the start of differentiation; 48 h later, AB678 cells were treated with ethidium bromide (EtBr) to inhibit nascent transcription of mt-mRNAs. Cells were collected at the indicated time points after adding EtBr to measure the time required for mt-mRNAs to reach one-half of the abundance measured at time 0 (the t_1/2_).^43–45^ As shown, silencing *LANCL1-AS1* generally reduced the stability of mt-mRNAs; for example, the t_1/2_ of *mt-ND1* mRNA declined from ∼3 h to ∼1.5 h, *mt-ND2* mRNA from ∼3.5 h to ∼2 h, and other mt-mRNAs followed a similar trend (Figure 6A). Notably, *mt-ND6* mRNA remained essentially unchanged after silencing *LANCL1-AS1*, in agreement with earlier results of steady-state mt-mRNA levels after silencing *LANCL1-AS1* (Figure 3C-E).^46^ In control experiments, a stable transcript, *GAPDH* mRNA (encoding the housekeeping protein GAPDH, transcribed in the nucleus by RNA polymerase II, which is refractory to EtBr), showed comparable stability whether *LANCL1-AS1* was silenced or not (Figure 6A). An additional control included was *MYC* mRNA, a short-lived transcript encoded by the nuclear genome and also transcribed by RNA polymerase II; as expected, the *MYC* mRNA relative stability was not affected by EtBr treatment, supporting reports that EtBr did not affect the stability of mRNAs transcribed in the nucleus.^44,45^

**Figure 6.**
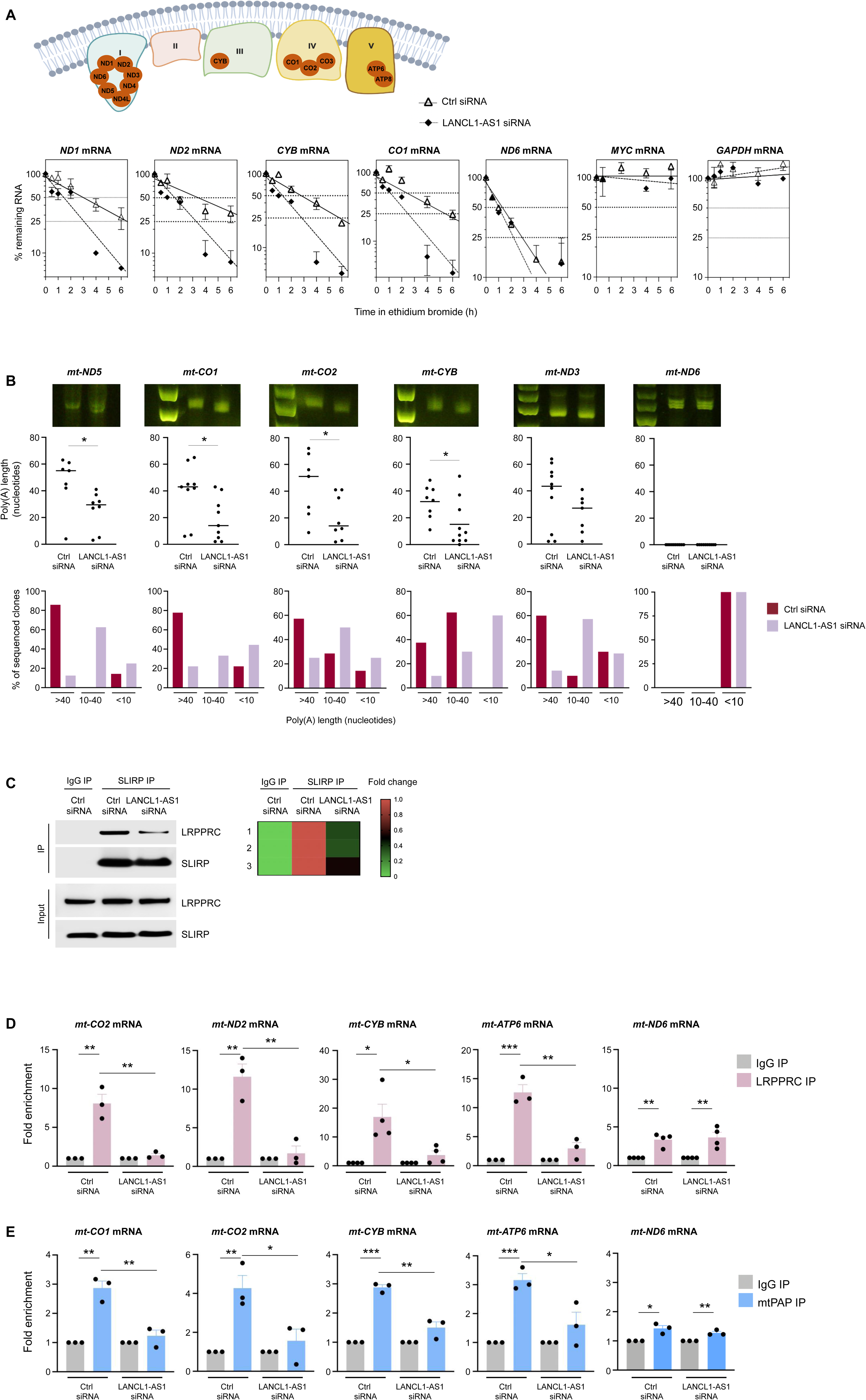
*LANCL1-AS1* increases poly(A) tail length of mt-mRNAs, promotes their stability. **(A)** *Top*, schematic depicting the protein components of the respiratory chain complexes I, III, IV and V, encoded by the mitochondrial genome; protein components of complex II are encoded by nuclear DNA. Created using BioRender. *Bottom*, 24 h after transfecting Ctrl siRNA or LANCL1-AS1 siRNA, AB678 myoblasts were placed in differentiation medium for an additional 72 h; cells were then treated with EtBr, and the relative levels of *mt-ND1*, *mt-ND2*, *mt-ND6*, *mt-CYB*, and *mt-CO1* mRNAs, as well as two control transcripts encoded by genomic DNA, *MYC* mRNA (unstable under conditions of RNA polymerase II inhibition), and *GAPDH* mRNA (a stable transcript), were assessed by RT-qPCR analysis and normalized to *18S* rRNA levels, which was also quantified by RT-qPCR analysis. The relative mRNA stabilities were assessed as the times required to reach 50% of the initial abundance of the mt-mRNAs at time 0 (before adding EtBr). **(B)** By 24 h after transfecting Ctrl or LANCL1-AS1 siRNAs, AB678 myoblasts were placed in differentiation medium for an additional 72 h. Cultures were then collected, and mt-mRNA poly(A) length was measured by the MPAT assay (STAR Methods); the poly(A) tail lengths of mt-mRNAs were compared between the Ctrl siRNA and LANCL1-AS1 siRNA transfection groups by analyzing changes in mobility of the bands spanning the poly(A) tails on agarose gels (*top*), and by measuring the length of poly(A) tails using Sanger sequencing after cloning the 3’ ends (*middle*), and by evaluating the relative proportion of mt-mRNAs bearing poly(A) tail lengths within different size ranges (*bottom*). **(C)** Cells that were processed as in (B) were further subjected to co-immunoprecipitation (co-IP) assays using an antibody that recognizes SLIRP or a control IgG. The levels of LRPPRC in the IP materials were then evaluated by western blot analysis; signals in ‘Input’ lysates, without IP, were also assessed (*left*). LRPPRC signals on western blots were quantified using Image J and represented as a heatmap (*right*). **(D,E)** By 24 h after transfecting Ctrl or LANCL1-AS1 siRNAs, AB678 myoblasts were differentiated for an additional 72 h; cells were then UV-crosslinked, and RIP assays (STAR Methods) using anti-LRPPRC relative to IgG antibodies (D) or anti-mtPAP relative to IgG antibodies (E) were performed in both transfection groups. The levels of LRPPRC-interacting mt-mRNAs were measured by RT-qPCR analysis. In D,E data are shown as the means ±SEM of three biological replicates. Significance (*, p<0.05; **, p<0.01; ***, p<0.001) was established using Student’s *t*-test.

Given that mt-mRNAs typically have no 3’UTR, the length of their poly(A) tails strongly influences their stabilities.^46–50^ We thus examined if *LANCL1-AS1* regulated the length of poly(A) tails of mt-mRNAs by carrying out poly(A) tail measurement assays. As shown, silencing *LANCL1-AS1* globally reduced the poly(A) tail lengths of mt-mRNAs, visualized here by comparing the sizes of specific PCR products generated to visualize the poly(A) tails of mt-mRNAs, and subjected to electrophoresis through agarose gels (Figure 6B, *top*). TA cloning followed by Sanger sequencing confirmed these results, namely that silencing *LANCL1-AS1* leads to higher levels of mt-mRNAs with short (<35-nt) poly(A) tails, suggesting that reduced levels of *LANCL1-AS1* globally reduced the length of poly(A) tails of mt-mRNAs (Figure 6B, *bottom*). Interestingly, when we measured *mt-ND6* mRNA, an mt-mRNA lacking a poly(A) tail^46,48^ (Figure 6B, *bottom*), the results showed no difference in mobility after electrophoresis through agarose gels (Figure 6B, *top*). Furthermore, *mt-ND6* mRNA showed no poly(A) tail by Sanger sequencing of AB678 cells, whether transfected with Ctrl or LANCL1-AS1 siRNAs, while other mt-mRNAs did (Figure 6B and Figure S5B). In sum, these results suggest that *LANCL1-AS1* promotes mitochondrial function at least in part by helping to maintain the lengths of poly(A) tails and stabilizing mt-mRNAs.

To elicit stabilization of mt-mRNAs, LRPPRC associates with another RBP, SLIRP, in an RNA-dependent manner.^51^ Moreover, SLIRP enhances the ability of LRPPRC to bind mt-mRNAs.^42,48,52^ Interestingly, ChIRP-western blot analysis suggested that *LANCL1-AS1* can interact with both LRPPRC and SLIRP (Figure 4C). Therefore, we studied how the lncRNP *LANCL1-AS1*-LRPPRC maintained mt-mRNA poly(A) tail length, and if *LANCL1-AS1* affected the formation of the LRPPRC-SLIRP chaperone complex, and thus modulated their binding to mt-mRNAs.

*LANCL1-AS1* was silenced in AB678 cells, and the interaction between LRPPRC and SLIRP was monitored by co-immunoprecipitation and western blot analysis. As shown, silencing *LANCL1-AS1* reduced the ability of SLIRP to interact with LRPPRC (Figure 6C and Figure S5C), suggesting that *LANCL1-AS1* may serve as a scaffold lncRNA to promote formation of this complex. To further test this hypothesis, we examined if loss of *LANCL1-AS1* reduced the binding capacity of LRPPRC to mt-mRNAs. After silencing *LANCL1-AS1* in AB678 cells and triggering differentiation for 48 h, UV-crosslinking followed by RIP assay revealed a significant reduction of LRPPRC binding to mt-mRNAs (Figure 6D). In keeping with our previous findings, *mt-ND6* mRNA interacted only modestly with LRPPRC, and this interaction showed no significant change after silencing *LANCL1-AS1* (Figure 6D). Moreover, LRPPRC maintains the poly(A) tail length of mt-mRNAs by recruiting the mitochondrial poly(A) polymerase (mtPAP, also known as PAPD1 and TUTase1);^46,49^ we thus investigated if silencing *LANCL1-AS1* might affect mtPAP binding to mt-mRNAs. RIP analysis revealed that mtPAP bound mt-mRNAs significantly in differentiated AB678 cultures. Importantly, silencing *LANCL1-AS1* for 48 h partially reduced these interactions (Figure 6E), but *mt-ND6* mRNA showed negligible binding to mtPAP and silencing *LANCL1-AS1* had no effect (Figure 6E). These observations indicate that *LANCL1-AS1* promotes polyadenylation of mt-mRNAs by facilitating the assembly of LRPPRC, SLIRP, and mtPAP on mt-mRNAs.

### Increasing *LANCL1-AS1* levels restores mitochondrial activity and myogenesis of old muscle progenitor cells

Mitochondrial activity declines with muscle aging, serving as a key pathological hallmark of sarcopenia.^53–57^ To assess the clinical relevance of our findings, we examined *LANCL1-AS1* expression levels in muscle from surgical hip replacement patients with sarcopenia or healthy controls (STAR Methods). Strikingly, *LANCL1-AS1* was consistently downregulated in sarcopenic muscle (Figure 7A), suggesting that its deficiency is closely linked to the mitochondrial decay observed during muscle wasting. The proteins encoded by the mitochondrial genome critically govern mitochondrial metabolism, although the regulatory mechanisms controlling their expression are not fully understood. The expression of these proteins is known to be tightly controlled via post-transcriptional mechanisms including length of poly(A) tails. In line with this concept and our findings, we analyzed the poly(A) tail length of mt-mRNAs in intact human muscle samples. We collected a total of 7 commercially available human skeletal muscle RNA samples, divided them into two groups, young (21-27 y.o.) and old (76-88 y.o.), and examined the poly(A) tail length of mt-mRNAs. As shown, mt-mRNA poly(A) tails were shorter in old compared with young skeletal muscle as monitored by gel mobility assays and Sanger sequencing (Figure 7B and Figure S5D). These findings suggest that the reduction in mt-mRNA poly(A) tail length could be a defining feature of muscle aging, and recapitulates the decline in mitochondrial function with advancing muscle age.^57–59^ They also lend support to the proposed functions of *LANCL1-AS1* and LRPPRC in mt-mRNA poly(A) length, stability, and mitochondrial function in intact muscle.

**Figure 7.**
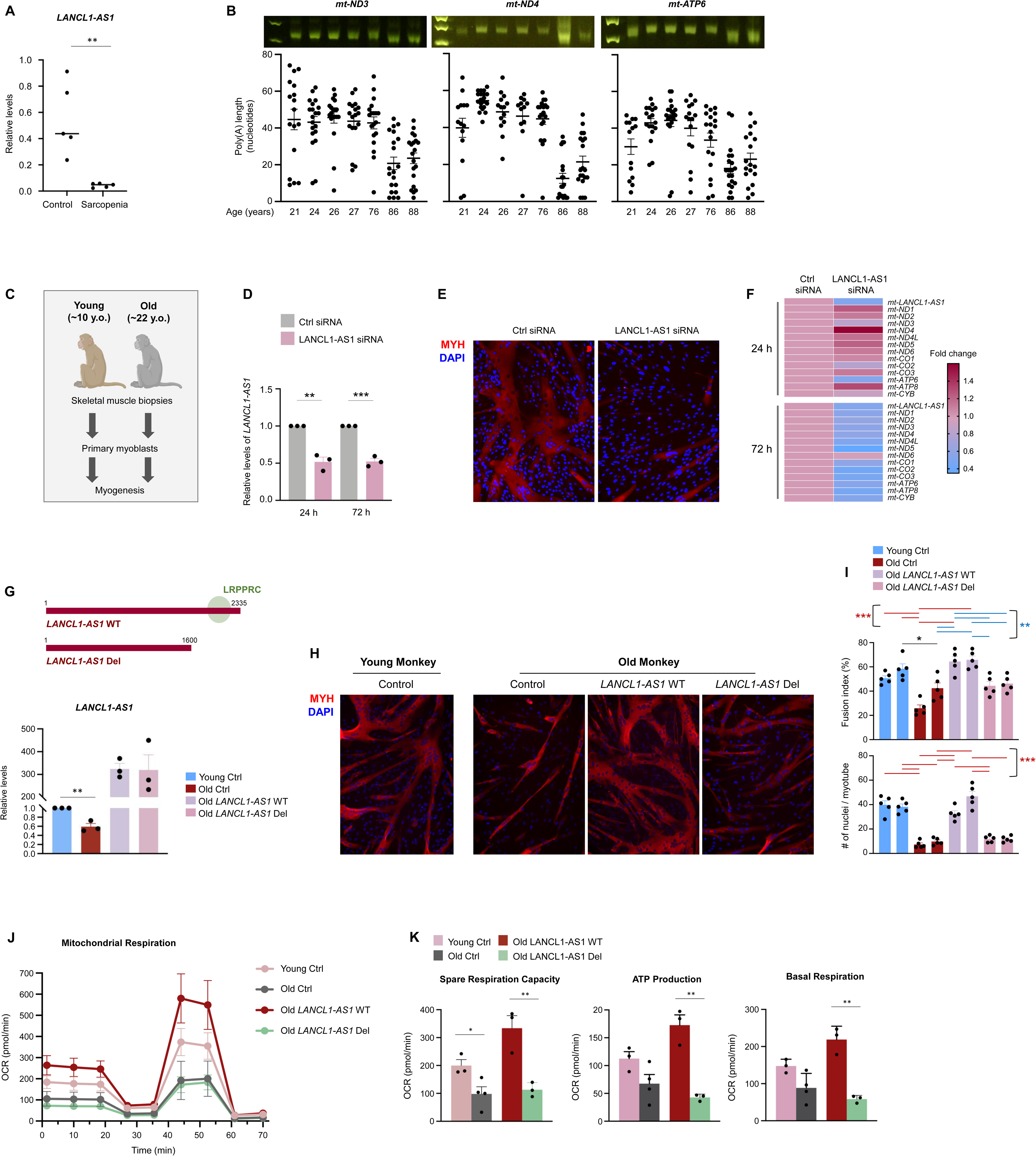
Poly(A) tail lengths of mt-mRNAs correlate with human muscle aging, while *LANCL1-AS1* supplementation restores myogenic capacity of primary myoblasts from old monkey skeletal muscle. **(A)** *LANCL1-AS1* levels in human skeletal muscle (gluteus maximus) biopsies harvested from healthy individuals (n=5) and patients with sarcopenia (n=5) were quantified using RT-qPCR analysis and normalized to the levels of *GAPDH* mRNA and *18S* rRNA. **(B)** MPAT assay (as described in Figure 6B) to compare the poly(A) tail lengths of mt-mRNAs in young and old human muscle biopsies; analysis was performed by comparing the mobility shifts of PCR products using DNA electrophoresis (*top*), and by precisely determining poly(A) tail lengths using Sanger sequencing after subcloning (*bottom*). **(C)** Skeletal muscle biopsies were collected from young (n=2) and old (n=2) rhesus monkeys and myoblasts were isolated for analysis. **(D-F)** Primary myoblasts from young monkeys were transfected with Ctrl or LANCL1-AS1 siRNAs; 24 h later, they were placed in differentiation medium for an additional 72 h, whereupon total RNA was extracted and the levels of remaining *LANCL1-AS1* measured by RT-qPCR analysis; data were normalized to *GAPDH* mRNA (D). The progression to differentiation into myotubes was monitored by immunofluorescent analysis to assess MYH levels (red) (E) and the levels of mt-mRNAs at 24 h and 72 h after inducing differentiation were calculated by RT-qPCR analysis and represented using a heatmap (F). **(G-I)** Schematic of the *LANCL1-AS1* transcripts expressed from viruses for this experiment: full length *LANCL1-AS1* WT, capable of binding LRPPRC, and a deletion mutant (*LANCL1-AS1* Del) lacking the distal 3’ end and hence unable to bind LRPPRC. RT-qPCR analysis was used to measure the expression levels of endogenous *LANCL1-AS1* in primary myoblasts from young and old monkey donors infected (3 MOI) with an empty vector (Ctrl), as well as the levels of *LANCL1-AS1* WT and *LANCL1-AS1* Del overexpressed in primary myoblasts from old monkey donors infected (3 MOI) with the respective vectors, 24 h after infection; *GAPDH* mRNA was used for normalization (G). Cultures processed as described in (G) were placed in myogenesis conditions; 72 h later, differentiation was compared between myoblasts from young and old monkey donors, as well as between old myoblasts overexpressing *LANCL1-AS1* WT relative to *LANCL1-AS1* Del by assessing MYH levels (red) detected by immunofluorescence (H) after staining nuclei with DAPI, and by calculating the fusion index and number of nuclei per myotube in five separate fields per experiment (I). **(J,K)** Mitochondrial activity was measured using a Seahorse instrument in the myoblast groups described in panel C to measure oxygen consumption rates (OCR) (J). Spare respiration capacity (*left*), ATP production (*middle*), and basal respiration (*right*) were calculated (K) (STAR Methods). Data in (B,D,G,I,K) represent the means ±SEM of three or more biological replicates. Significance (*, p<0.05; **, p<0.01; ***, p<0.001) was established using Student’s *t*-test.

*LANCL1-AS1* is a primate-specific lncRNA, with a high degree of sequence conservation between human and monkey, but not mouse (Figure S5E). Given the lack of conservation of *LANCL1-AS1* in mice, to study its physiologic role *in vivo*, we adopted instead a non-human primate *ex vivo* model to explore *LANCL1-AS1* function. Specifically, we investigated the impact of *LANCL1-AS1* in muscle aging in rhesus monkeys (*Macaca mulatta*) from the National Institute on Aging (NIA) by collecting muscle biopsies from young (∼ 10 years old, y.o.) and old (∼ 22 y.o.) monkeys and preparing primary myoblasts to study myogenesis (STAR Methods; Figure 7C). *LANCL1-AS1* was silenced in young monkey myoblasts at 24 and 72 h after transfection of species-specific siRNAs (Figure 7D). Twenty-four h after transfecting *LANCL1-AS1* siRNAs, myoblasts were placed in differentiation conditions; as shown, myogenesis was potently reduced relative to Ctrl siRNA (Figure 7E), as determined by visualizing myofibers using MYH fluorescence 72 h later. Along with reducing myogenesis, RT-qPCR analysis revealed that silencing *LANCL1-AS1* lowered the production of most monkey mt-mRNAs late in differentiation (72 h), but not *mt-ND6* mRNA (Figure 7F). Early in differentiation (24 h), silencing *LANCL1-AS1* reduced the levels of only a few mt-mRNAs such as *mt-ND3*, *mt-CO2*, and *mt-ATP6* mRNAs (Figure 7F).

In monkey primary myoblasts, we were further able to study the effect of restoring *LANCL1-AS1* function on myogenesis. We prepared lentiviral vectors to express human *LANCL1-AS1* that was either full-length (wild-type, WT) or truncated by deletion of the 3’ end (*LANCL1-AS1* Del) and hence unable to bind LRPPRC (Figure 7G). In agreement with human and mouse primary myoblasts,^53–55^ young monkey myoblasts displayed higher myogenic capacity than old monkey myoblasts, as determined by monitoring myotube formation and by quantifying the fusion index and number of nuclei per myotubes at 72 h into the myogenic program (Figure 7H and 7I). Strikingly, lentivirus-driven ectopic expression of *LANCL1-AS1* WT largely restored the myogenic capacity of old myoblasts, with improved fusion indices and numbers of nuclei per myotube, while overexpressing *LANCL1-AS1* Del did not (Figure 7H and 7I).

In light of the fact that muscle mitochondrial activity declines with aging,^57–59^ we asked if the re-expression of *LANCL1-AS1* WT in old myoblasts might improve mitochondrial activity as it promotes myogenic regeneration. Young myoblasts, which expressed higher levels of *LANCL1-AS1*, displayed higher mitochondrial activity than old myoblasts, as assessed by monitoring OCR (including spare capacity, ATP production, and basal respiration; STAR Methods), which expressed lower *LANCL1-AS1* and displayed lower OCR (Figure 7J and 7K). Ectopic overexpression of *LANCL1-AS1* WT restored mitochondrial activity of old monkey myoblasts to levels even superior to those of young myoblasts, while overexpressing *LANCL1-AS1* Del did not improve mitochondrial activity above that of old myoblasts (Figure 7J and 7K). These results reflect key similarities in differentiating primary monkey myoblasts and primary human myoblasts (Figure 2), including the patterns of expression of mt-mRNAs and myogenic differentiation and fusogenic activities.^53^

## DISCUSSION

The loss of skeletal muscle with aging, sarcopenia, critically contributes to frailty, metabolic decline, and loss of mobility and independence.^57–59^ Here, a comparison of the transcriptomes of aging human skeletal muscle and human myogenesis revealed that *LANCL1-AS1* was among the most robustly declining lncRNAs in aging muscle, and most markedly increased lncRNAs during human myogenic differentiation. We further discovered that *LANCL1-AS1* forms a lncRNP complex with LRPPRC and modulates poly(A) tail length in mt-mRNAs, in turn promoting mitochondrial function and myogenic capacity. We propose that *LANCL1-AS1* contributes to maintaining mitochondria homeostasis in young skeletal muscle, and its loss in old muscle contributes to mitochondrial dysfunction and reduced myogenic capacity (Figure 8).

**Figure 8.**
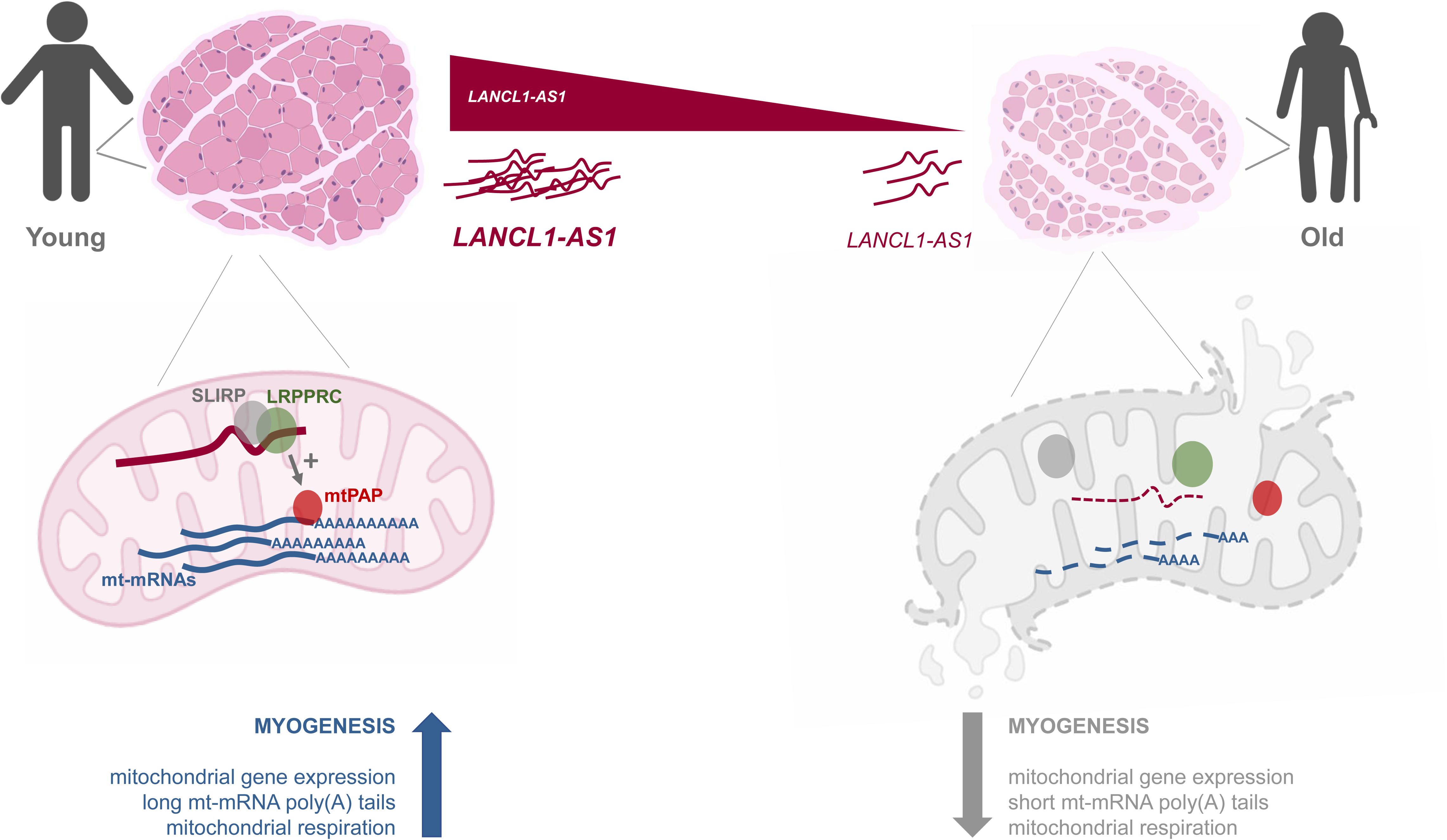
Proposed model. In young human skeletal muscle, the presence of abundant *LANCL1-AS1* promotes the function of LRPPRC-SLIRP, in turn helping to maintain mt-mRNA poly(A) tail lengths, which preserves mitochondrial homeostasis and promotes myogenesis. In old human skeletal muscle, *LANCL1-AS1* levels decline, leading to reduced function of the LRPPRC-SLIRP complex, shortened mt-mRNA poly(A) tails, and attenuated mitochondrial activity and myogenesis.

Even though *LANCL1-AS1* has been detected in other tissues, such as lung,^60,61^ a brief survey of cultured cell types revealed that *LANCL1-AS1* was highly expressed in skeletal muscle (Figure 1G), displaying a tissue-selective expression pattern, as do other lncRNAs; for example, *lincMD1* is only highly expressed in mouse skeletal muscle but not in other tissues like liver or heart.^16^ *LANCL1-AS1* appeared to be expressed in cultured neuroblastoma cells (Figure 1G), but its possible influence on processes like neurogenesis has not been reported. Interestingly, the protein expressed from the locus (LANCL1) is highly expressed in neuronal tissues, is found across the cytoskeleton and plasma membrane, and functions as a glutathione transferase, reducing neuronal oxidative stress.^62^ Whether LANCL1 protein functions similarly in skeletal muscle also remains to be studied. An intriguing possibility is that the function of *LANCL1-AS1*, which promotes mitochondrial gene expression and mitochondrial activity, and could therefore increase oxidative damage, might be balanced by the LANCL1 protein, which lowers oxidative stress. This hypothesis, as well as the possibility that LANCL1 is expressed in muscle, awaits testing. Additionally, although *LANCL1-AS1* was found in lung cancer,^60,61^ a more comprehensive survey of *LANCL1-AS1* expression across malignancies is needed. It will similarly be important to study if *LANCL1-AS1* production in cancer cells confers a tumorigenic advantage by increasing mitochondrial function.

*LANCL1-AS1* exhibits a species-specific expression pattern, with robust expression in human and primate skeletal muscle cells, but no expression in mouse skeletal muscle (not shown). As mentioned earlier, a lack of conservation across species is not unique to *LANCL1-AS1*, as many other lncRNAs fail to be conserved at the nucleotide level, even if specific functional domains of lncRNAs may be conserved, and some lncRNAs unrelated in their primary sequence have evolved to retain segments with orthologous functions across species.^63,64^ A mouse ortholog of *LANCL1-AS1* has not been identified, which limited our ability to carry out studies in murine model organisms. However, *LANCL1-AS1* is conserved in non-human primates, which allowed us to perform additional mechanistic experiments in Rhesus macaques (Figure 7).

A context-specific expression pattern in human skeletal muscle was also identified for *LANCL1-AS1*, which showed high levels of expression in young tissues, and declining abundance with advancing age. The transcriptional and post-transcriptional regulatory factors that govern *LANCL1-AS1* expression levels in muscle have not yet been identified. As we evaluate *LANCL1-AS1* as a possible target of intervention in muscle regeneration, it will be important to elucidate which transcription factors and post-transcriptional regulators (e.g., microRNAs, RBPs, etc.) govern *LANCL1-AS1* levels during myogenic regeneration to maintain skeletal muscle health.

In addition to changing abundance, the subcellular localization of *LANCL1-AS1* also appeared to be regulated, with mobilization from the nucleus to mitochondria as myogenesis progressed. This change in subcellular abundance is reminiscent of the translocation of the cytoplasmic lncRNA *SAMMSON*, whose expression levels were highly increased in melanoma cells.^65^ The tumor-promoting function of *SAMMSON* was associated with its ability to enhance mitochondrial activity in melanoma, as loss of *SAMMSON* disrupted mitochondrial functions and suppressed melanoma progression.^65^ Similarly, *LANCL1-AS1* appeared to translocate from the nucleus to the cytoplasm and mitochondria as differentiation advanced (Figure 4J, Figure 5A and 5B). The specific mechanisms that mediate *LANCL1-AS1* export to the cytoplasm and further mobilize *LANCL1-AS1* into mitochondria are not fully understood at the moment. We identified 20% of *LANCL1-AS1* as being present in the mitochondria (Figure 5D and 5E), and found evidence that PNPT1 (PNPase) partially contributes to the import of *LANCL1-AS1* into mitochondria (Figure 5G and 5H). Whether additional transport factors, for example VDAC^66^ are responsible for the mitochondrial internalization of *LANCL1-AS1* remains to be studied.

In the mitochondria, *LANCL1-AS1* was found to promote the polyadenylation of mt-mRNAs, and thereby enhanced mitochondrial respiration to generate energy for muscle differentiation and function. We found that *LANCL1-AS1* elicited this function by interacting with the mitochondrial protein LRPPRC (Figures 4 and 5). We further uncovered that silencing *LANCL1-AS1* reduces the binding of LRPPRC and SLIRP (Figure 6C). The fact that the LRPPRC-SLIRP complex promotes mtPAP activity,^39,40^ helps explain why silencing *LANCL1-AS1* effectively reduced polyadenylation of mt-mRNAs, lowering their stability and production of encoded respiratory chain proteins (Figure 3G, Figure 6A-C). The discovery that *LANCL1-AS1* is required for mt-mRNA polyadenylation and stabilization, leading to increased respiration, warrants further consideration. Perhaps *LANCL1-AS1* is a marker of muscle function robustness, or perhaps it is involved in other systems that rely on high levels of energy, like the nervous system. Further support for this future work comes from evidence that mutations in LRPPRC cause Leigh syndrome French Canadian variant (LSFC), a neurodegenerative disease linked to an impairment in energy homeostasis.^42,67^

Dysregulated mitochondrial activity is a widespread feature of aging tissues^68^ and influences age-associated physiologic declines and diseases, including cardiovascular pathologies, neurodegeneration, cancer, and sarcopenia—supporting the long-standing view that mitochondria homeostasis is important for health and longevity. Here we have found that *LANCL1-AS1* is linked to muscle aging, myogenesis, and the maintenance of mt-mRNA poly(A) length and mitochondrial activity (Figure 8). It will be important to investigate if mt-mRNA poly(A) tail lengths and/or *LANCL1-AS1* levels inform about mitochondrial function and organ aging, particularly as we seek markers to track the effects of aging and improve overall health and longevity.

### Limitations of the study

An important gap in this work is our incomplete knowledge at present of the subcellular localization of *LANCL1-AS1* in myoblasts and myotubes. Similarly, the specific mitochondrial region where *LANCL1-AS1* resides has not been elucidated; high-definition confocal or electron microscopy are needed to establish its localization. The full set of factors contributing to the import of *LANCL1-AS1* into mitochondria, working alongside or perhaps independently of PNPT1, also remains to be identified.

The stoichiometry of *LANCL1-AS1* relative to mitochondria in this system also remains to be fully understood. Despite finding ∼75–115 copies of *LANCL1-AS1* per nucleus equivalent in myotubes, the number of mitochondria likely exceeds that number by tenfold or more. Whether *LANCL1-AS1* is only employed to supplement constitutive polyadenylation of mt-mRNAs in a subset of mitochondria at specific times also remains to be further studied, as tools for single-mitochondria lncRNP analysis become available.

A key limitation of this study is that our experimental system primarily evaluates myogenic differentiation and mitochondrial function in cultured myoblasts rather than the full process of skeletal muscle regeneration *in vivo*. Although myoblast differentiation is an essential component of regeneration, successful regeneration additionally depends on satellite cell activation and self-renewal, inflammatory responses, fibro-adipogenic progenitors, vascular cells, extracellular matrix remodeling, and other processes occurring in the muscle niche. Therefore, our findings should be interpreted as identifying a role for *LANCL1-AS1* in myogenic capacity rather than establishing a direct role in whole-muscle regeneration. Likewise, while reduced *LANCL1-AS1* expression in old and sarcopenic muscle is consistent with a potential contribution to age-related muscle dysfunction, sarcopenia is a condition influenced by many mechanisms beyond regenerative decline, such as denervation, endocrine and metabolic changes, chronic inflammation, and mitochondrial dysfunction. Our data support a model in which loss of *LANCL1-AS1* contributes to impaired mitochondrial function and diminished myogenic differentiation in aging muscle cells; but the extent to which it influences muscle regeneration or sarcopenia *in vivo* remains to be determined.

A major impediment to understanding more fully the *in vivo* role of *LANCL1-AS1* in health and disease is the inability to develop a mouse model, as *LANCL1-AS1* is not conserved in rodents. If a functional ortholog of the human *LANCL1-AS1* is identified in mouse, experiments to delete it and overexpress it across the body and in a tissue-specific manner will be particularly informative. Given that the activation of muscle stem cells also requires a shift in metabolic utilization to increase mitochondrial activities, other important experiments on a possible ortholog should be aimed at determining if *LANCL1-AS1* plays a role in the activation of muscle stem cells during muscle regeneration. Experiments are underway to overexpress *LANCL1-AS1* in mouse skeletal muscle to study muscle regeneration and to identify a mouse ortholog with a similar function, so that we can understand the impact of *LANCL1-AS1* in physiologic context.

---

**Dataset 1. LncRNAs differentially abundant in skeletal muscle as a function of age in healthy individuals.** Long noncoding (lnc)RNAs [including antisense lncRNAs and long intergenic noncoding RNAs (lincRNAs)] identified as changing significantly (adjusted p-value<0.05) in skeletal muscle biopsies from healthy human participants in the GESTALT study (92 participants total), spanning ages from 22 to 89 (STAR Methods).

**Dataset 2. LncRNAs differentially abundant during human myogenesis in culture.** RNA-seq analysis of human skeletal myoblasts differentiating in culture [including previously reported RNA-seq datasets in GSE215343, as well as GSE261476 (token oterkuicznajdij)]. Six lncRNAs shared with the lncRNAs identified in the GESTALT study (Dataset 1) are shown in Figure 1C.

## STAR * METHODS

Detailed methods are provided in the online version of this paper and include the following:

- KEY RESOURCES TABLE
- RESOURCE AVAILABILITY

- Lead contact
- Materials availability
- Data and code availability
- EXPERIMENTAL MODEL

- Cell culture and myogenic differentiation
- Criteria for inclusion of GESTALT participants
- Isolation of primary monkey fibroblasts
- Human muscle biopsies
- METHOD DETAILS

- Creatine kinase activity
- 5’RACE
- Reverse transcription (RT) and real-time quantitative (q)PCR analysis
- Western blot analysis
- Immunocytochemistry and Confocal Imaging
- Library preparation protocol
- RNA-seq analysis
- Modified ChIRP analysis for mass spectrometry (MS) and western blot analyses of RNA-bound proteins
- Mass spectrometry
- smFISH
- Mitochondrial poly(A) tail (MPAT) assays
- Mitochondrial activity measurement using Seahorse
- QUANTIFICATION AND STATISTICAL ANALYSIS

## ACKNOWLEGMENTS

We thank the staff in the Department of Orthopaedics Clinics and Traumatology and Navarrabiomed Biobank for their support during the study, as well as our patients and their families. We are grateful to the GESTALT participants and the GESTALT Study Team at Harbor Hospital (Baltimore, MD), and NIA for their contribution to sample collection and project coordination.

## AUTHOR CONTRIBUTIONS

J.H.Y designed the project. J.H.Y., E.K.I., K.M.M., D.T., B.R., C.S., R.M., J.L.M., C.A., M.S., M.R., Y.P., J.F., Y.C.C., B.A.C.V., R.F., T.H.C., S.D., and P.S performed experiments and analyzed data. J.H.Y., M.B., N.M.V., M.M., and M.G. supervised and coordinated the study. J.A.M, R.dC., L.F., X.Y., K.A., and C.Y.C. provided reagents and intellectual contributions. J.H.Y. and M.G. wrote the paper, with input from all authors.

## INCLUSION AND DIVERSITY

We support inclusive, diverse, and equitable conduct of research. One or more of the authors of this paper self-identifies as an underrepresented ethnic minority in their field of research or within their geographical location. One or more of the authors of this paper self-identifies as a gender minority in their field of research.

## STAR * METHODS

### KEY RESOURCES TABLE

| REAGENT or RESOURCE | SOURCE | IDENTIFIER |
| --- | --- | --- |
| **Antibodies** |  |  |
| MYH/MHC (B-5) | Santa Cruz Biotechnology | sc-376157 |
| HSP90 (F-8) | Santa Cruz Biotechnology | sc-13119 |
| MEF2C (D80C1) | Cell Signaling Technology | 5030 |
| Total OXPHOS Human WB Antibody Cocktail including ATP5A, UQCRC2, SDHB, COX2, NDUFB8 | Abcam | ab110411 |
| SLIRP | Abcam | ab51523 |
| LRPPRC (F-7) | Santa Cruz Biotechnology | sc-166178 |
| HADHB (E-1) | Santa Cruz Biotechnology | sc-271495 |
| mtPAP (1D3) | GeneTex | GTX70156 |
| VDAC (D73D12) | Cell Signaling Technology | 4661 |
| 7 commercially available human skeletal muscle RNA samples: young (21-27 y.o.) and old (76-88 y.o.) | BioChain | Cat# R1234171-50 |
| **Bacterial and virus strains** |  |  |
| Empty Lentivirus pLenti-P2A-tGFP | Origene | Cat# PS100108 |
| Lenti-LANCL1-AS1 (WT) | This study |  |
| Lenti-LANCL1-AS1 (Del) | This study |  |
| **Critical commercial assays** |  |  |
| Poly(A) Tail-Length Assay Kit | Invitrogen | Cat# 764551KT |
| Direct-zol RNA MiniPrep Kit | Zymo Research | Cat# R2052 |
| TOPO TA Cloning Kit | Invitrogen | Cat# 450030 |
| QIAquick PCR Purification Kit | Qiagen | Cat# 28104 |
| EnzyChrom Creatine Kinase Assay Kit | Bioassay Systems | Cat# ECPK-100 |
| Ovation® RNA-Seq System V2 | Nugen | Cat# 7102 |
| TruSeq ChIP Library Preparation Kit | Illumina | Cat# IP-202-1012 |
| Low Input RiboMinus Eukaryote System v2 | Thermo Scientific | Cat# A15027 |
| 5'/3' RACE Kit, 2<sup>nd</sup> generation | Sigma | Cat# 3353621001 |
| XF Cell Mito Stress Kit | Agilent Technologies | Cat# 103015-100 |
| **Chemicals, peptides, and recombinant proteins** |  |  |
| Lipofectamine 3000 | Invitrogen | Cat# L3000150 |
| Lipofectamine RNAiMAX | Invitrogen | Cat# 13778150 |
| Hyclone Ham's F10 media | Cytiva | Cat# SH30025.01 |
| Skeletal Muscle Cell Growth Media | Promocell | Cat# C-23060 |
| DMEM | Gibco | Cat# 11965092 |
| FBS | Gibco | Cat# A5256801 |
| Horse Serum | Gibco | Cat# 16050114 |
| Penicillin-Streptomycin | Gibco | Cat# 15140122 |
| Nonessential Amino Acids (100X) | Gibco | Cat# 11140050 |
| Antibiotic-antimycotic (100X) | Gibco | Cat# 15240062 |
| Gentamicin solution | Sigma | Cat# G1272 |
| Plasmocin® prophylactic | InvivoGen | Cat# ant-mpp |
| Dispase II protease | Sigma | Cat# D4693 |
| Dispase II | Gibco | Cat# 17105041 |
| Chicken Embryo Extract | MP Biomedicals | Cat# 092850145 |
| Collagenase D | Roche | Cat # 11088858001 |
| Collagenase, Type II, powder | Gibco | Cat# 17101015 |
| Matrigel Basement Membrane Matrix | Corning | Cat# 356235 |
| BioCoat Matrigel Matrix Cellware, Corning - 6-Well (Thin Layer) | Corning | Cat# 354603 |
| Corning BioCoat Cellware, Collagen Type I, Corning-6-Well | Corning | Cat# 354400 |
| bFGF | ORF Genetics (Iceland) | Cat# 01-A1110-0100 |
| Normal Goat Serum (10%) | Thermo Scientific | Cat# 50062Z |
| Triton X-100 | Sigma | Cat# X100 |
| Paraformaldehyde (16%) | Electron Microscopy Sciences | Cat# 15710 |
| Halt Phosphatase Inhibitor | Thermo Scientific | Cat# 78420 |
| Proteinase K | Invitrogen | Cat# 25530049 |
| Halt Protease Inhibitor | Thermo Scientific | Cat# 87786 |
| Maxima Reverse Transcriptase | Thermo Scientific | Cat# EP0743 |
| TriPure Isolation Reagent | Roche | Cat# 11667165001 |
| KAPA SYBR Fast ABI Master Mix (2X) | Roche | Cat# 07959435001 |
| Complete Protease Inhibitor Cocktail (EDTA-free) | Roche | Cat# 11873580001 |
| RiboLock RNase Inhibitor | Invitrogen | Cat# EO0381 |
| Random Primers | Invitrogen | Cat# 48190011 |
| dNTP Mix (10 mM each) | Thermo Scientific | Cat# R0192 |
| **Deposited data** |  |  |
| KM155 RNA-seq | GEO | GSE261476<br>(reviewer token oterkuicznajdij) |
| AB678 si-LANCL1-AS1 RNA-seq | GEO | GSE261477<br>(reviewer token yzqxkuykfvpbqx) |
| **Experimental models: Cell lines** |  |  |
| AB678 | V. Mouly | doi:<br>10.1186/s13395-016-0115-5 |
| AB1167 | V. Mouly | doi:<br>10.1186/s13395-016-0115-5 |
| Monkey young and old primary myoblasts | This study | NIA |
| KM155 | V. Mouly | doi:<br>10.1186/s13395-016-0115-5 |
| **Oligonucleotides** |  |  |
| RT-qPCR primers, see Table S1 | This study | N/A |
| ChIRP-MS ASO, see Table S2 | This study | N/A |
| siRNA for silencing LANCL1-AS1, LRPPRC, see Table S3 | This study | N/A |
| MPAT primers, see Table 4 | This study | N/A |
| MS data by ChIRP-MS, see Table 5 | This study | N/A |
| smFISH probes, see Table S5 | This study | N/A |
| **Software and Algorithms** |  |  |
| bcl2fastq (v2.20.0.422) | Illumina | https://support.illumina.com/sequencing/sequencing_software/bcl2fastq-conversion-software.html |
| cutadapt (v1.18) | Martin, 2011 | https://cutadapt.readthedocs.io/en/v1.18/ |
| STAR (v2.4.0j) | Dobin et al., 2013 | https://github.com/alexdobin/STAR |
| featureCounts (v1.6.4) | Liao et al., 2014 | https://subread.sourceforge.net/ |
| DESeq2 (v1.26.0 & v1.30.0) | Love et al., 2014 | https://bioconductor.org/packages/release/bioc/html/DESeq2.html |
| clusterProfiler (v4.2.2) | Yu et al., 2012 | https://guangchuangyu.github.io/software/clusterProfiler/ |
| Python | Python | https://www.python.org/ |
| R Studio | R Studio | https://www.r-project.org/ |
| **others** |  |  |
| Verity 96-well Thermal cycler | Applied Biosystems | Cat# 4375305 |
| QuantStudio 5 Real-Time PCR System | Applied Biosystems | Cat# A28140 |
| Agilent Technologies 2100 Bioanalyzer | Agilent Technologies | Cat# G2939A |
| Illumina HiSeq 2500 Sequencing System | Illumina Inc. | Cat# SY-401-2501 |
| Keyence BZX710 All-in-One Fluorescence microscope | Keyence Corporation | Cat# BZX710 |
| SeaHorse XFe24 Flux Analyzer | Agilent Technologies | Model XFE24 |

### RESOURCE AVAILABILITY

#### Lead contact

Further information and requests for resources and reagents should be directed to and will be fulfilled by the lead contact, Myriam Gorospe.

#### Materials availability

All unique/stable reagents generated in this study are available from the lead contact with a completed Materials Transfer Agreement.

#### Data and code availability

Sequencing data have been deposited (GSE261476, reviewer token oterkuicznajdij; GSE261477, reviewer token yzqxkuykfvopbqx). Sequencing data previously published that was analyzed here were obtained from GSE164471. Access to GESTALT data is available upon review and subsequent approval of proposals submitted through the BLSA study website (https://www.blsa.nih.gov/). Other original data reported in this paper will be shared by the lead contact upon request. This paper does not report original code.

### EXPERIMENTAL MODEL

#### Cell culture and myogenic differentiation

Immortalized human AB1167, AB678, and KM155 myoblasts were developed and cultured as described.^21,28^ Briefly, human myoblasts were cultured in growth medium [equal volume mixture of Ham’s F10 medium supplemented with 20% fetal bovine serum (FBS) and Promocell skeletal muscle cell growth medium] and were induced to differentiate by growth to high density and replacement of the growth medium with differentiation medium [Dulbecco’s modified Eagle medium (DMEM) supplemented with 2% horse serum]. For silencing experiments, using a final concentration of 50 nM siRNA and Lipofectamine RNAiMAX (Thermo Fisher Scientific), control small interfering RNA (Ctrl siRNA), LANCL1-AS1 siRNA, LRPPRC siRNA, or PNPase/PTPN1 siRNA was transfected 24 h before induction of differentiation (Table S2). For overexpression of full-length, wild type *LANCL1-AS1* (*LANCL1-AS1 WT*) or a truncated version with a 3’ deletion that prevented binding of LRPPRC (*LANCL1-AS1 Del*), lentiviruses were constructed and used at a final concentration of 3 MOI (multiplicity of infection).

#### Criteria for inclusion of GESTALT participants

Participants in the GESTALT study, a longitudinal cohort of healthy adults aged 20 years and older were recruited from the Washington, DC, and Baltimore, MD areas. Eligibility required willingness to return for follow-up evaluations every two years, consent for collection and long-term storage of DNA and RNA samples, body mass index (BMI) <30 kg/m², eligibility for magnetic resonance imaging (MRI), and absence of established genetic or autoimmune diseases. Participants were selected using stringent criteria designed to identify individuals with ideal health status across the adult lifespan.^69^ Eligible participants had no history of cardiovascular, pulmonary, gastrointestinal, metabolic, neurological, kidney, liver, or autoimmune diseases; no active cancer or cancer within the previous 10 years (except limited non-melanoma skin cancer or successfully treated cancers without recurrence for at least 10 years); no cognitive or functional impairment; and no major psychiatric disorders. Participants were required to demonstrate preserved physical function, including independence in activities of daily living and high performance on standardized physical and cognitive assessments. Additional eligibility criteria included normal hematologic and biochemical laboratory values and absence of chronic use of medications indicative of significant underlying disease. Once enrolled, participants remained in the cohort even if their health status changed during follow-up.

#### Isolation of primary monkey myoblasts

Muscle progenitor cells were isolated from four rhesus monkeys (*Macaca mulatta*; 2 young and 2 old, as shown below) according to a protocol approved by the Intramural Research Program’s ACUC. Briefly, monkey gastrocnemius wedge muscle biopsies were minced and enzymatically digested using digestion solution (PBS with 500 U/mL collagenase II, 2.5 U/mL dispase II, 1.5 U/mL collagenase D, and 2.5 mM CaCl_2_). The muscle tissue fragments were seeded on tissue culture plates coated with 0.9 mg/mL Matrigel to allow the outgrowth of muscle progenitor cells from the muscle explants. The expanded single cells were collected and detached using trypsin and transferred to a plate coated with collagen (0.01%) for 1 h at 37°C to remove epithelial cells and fibroblasts. One hour later, floating cells (containing the muscle progenitor cells) were seeded on an uncoated 12-well plate in growth medium composed of high-glucose DMEM, 20% fetal bovine serum, 10% horse serum, 0.5% chicken embryo extract, 2.5 ng/mL bFGF, 10 mg/mL Gentamycin, 1% Antibiotic-Antimitotic, 2.5 µg/mL Plasmocin prophylactic.

| ID | Sex | Age (y.o.) | Biopsy Date | Comments |
| --- | --- | --- | --- | --- |
| Y1 | F | 10.1 | 06/29/22 | Gastrocnemius wedge |
| Y2 | M | 10.9 | 06/29/22 | Gastrocnemius wedge |
| O1 | M | 21.9 | 06/29/22 | Gastrocnemius wedge |
| O2 | M | 22.2 | 06/29/22 | Gastrocnemius wedge |

##### Human muscle biopsies

Muscle (gluteus maximus) was harvested from hip replacement surgeries in age-matched individuals who had normal muscle (Control, n=5) or had Sarcopenia (n=5). The control group consisted of five participants (3 males and 2 females) with a mean age of 78.6 years (range, 71–87 years), while the sarcopenia group consisted of five female participants with a mean age of 83.6 years (range, 78–90 years). Sarcopenia patients follow the 4 criteria below, none of the control patients followed any of the 4 criteria. Due to the acute conditions, perfomance measures were not taken into account for sarcopenia diagnosis.

| Consensus/Study | ASM (kg) | ALM/BMI (kg/BMI) | ALM/ht<sup>2</sup> (kg/m<sup>2</sup>) | Handgrip Strength (kg) |
| --- | --- | --- | --- | --- |
| **EWGSOP2 (2019)** DOI: 10.1093/ageing/afy169 | Men < 20 kg<br>Women < 15 kg | — | Men < 7.0 kg/m<sup>2</sup><br>Women < 5.5 kg/m<sup>2</sup> | Men < 27 kg<br>Women < 16 kg |
| **FNIH (2014)** DOI: 10.1093/gerona/glu010 | Men < 19.75 kg<br>Women < 15.02 kg | Men < 0.789<br>Women < 0.512 | — | Men < 26 kg,<br>Women < 16 kg |
| **IWGS (2011)** DOI: 10.1016/j.jamda.2011.01.003 | — | — | Men ≤ 7.23 kg/m<sup>2</sup><br>Women ≤ 5.67 kg/m<sup>2</sup> | — |
| **SCWD (2011)** DOI: 10.1016/j.jamda.2011.04.014 | — | — | Men ≤ 6.81 kg/m<sup>2</sup><br>Women ≤ 5.18 kg/m<sup>2</sup> | — |

The study followed the principles of the Declaration of Helsinki and was approved by the Navarra Research Ethics Committee (PI_2020/125), Spain.

### METHOD DETAILS

#### Creatine kinase activity

Creatine kinase (CK) activity was determined in cell lysates by using the EnzyChrom creatine kinase assay kit (BioAssay Systems) following the manufacturer’s protocol. Briefly, cell lysates (1 μg) were incubated with 10 µL substrate solution, 100 µL assay buffer and 1 µL enzyme mix at 37 °C for 20 min; reactions were read 20 and 40 min later at 340 nm. CK activity was calculated by the equation CK = (OD_40min_ – OD_20min_ /OD_CALIBRATOR_ – OD_H2O_) × 150, and expressed as ‘units per μg of total protein’ or ‘fold change’.

#### 5’RACE

The 5’ rapid amplification of cDNA ends (RACE) was performed using the 5’/3’ RACE Kit, 2nd Generation (Sigma) following the manufacturer’s instructions. The primer sequences for 5’ RACE of *LANCL1-AS1* were CAGGATTTGGGACAGGATTAGG (Sp1); GGGACAGGATTAGGTTCAGTTATC (Sp2); GAAGGCCAGAAAGAGTTGGT (Sp3).

#### Reverse transcription (RT) and real-time quantitative (q)PCR analysis

Total RNA from cultured cells was isolated using the Direct-zol^TM^ RNA MiniPrep kit (Zymo Research), which includes a digestion step using DNase I. For cDNA synthesis, reverse transcription (RT) was performed for RNA prepared in TriPure Isolation Reagent (Roche) using Maxima Reverse Transcriptase (Thermo Fisher Scientific) following the manufacturer’s protocol. Quantitative (q)PCR analysis of the cDNA was performed following the manufacturer’s instructions for KAPA SYBR FAST ABI Prism qPCR kit (KAPA Biosystems) with mRNA-specific primers (Table S3). RT-qPCR reactions were performed on QuantStudio 5 Real-Time PCR System (Thermo Fisher Scientific) with a cycle setup of 2 min at 95 °C and 40 cycles of 5 sec at 95 °C plus 20 sec at 60 °C; the fold change in abundance was calculated as described previously.^21,29^ In qPCR amplification reactions, control ‘RT minus’ (‘RT-’) reactions were routinely included. *LANCL1-AS1* copy numbers per cell in myoblasts and nucleus-equivalent in myotubes were calculated by using droplet digital PCR analysis, by interpolating on a standard curve of known nucleic acid concentrations, and by comparing relative Ct values of *OIP5-AS1* with the Ct values of a transcript of known abundance, *GAPDH* mRNA (estimated to be present at ∼1300 copies per cell and nucleus-equivalent, as previously determined).^22,23,29^

Total RNA was extracted from ∼200 mg of human muscle biopsies (gluteus maximus) that had been preserved in RNAlater Ice (Zymo Research). Tissue homogenization was performed in a bead mill homogenizer (30 Hz) using a 6-mm steel bead and RNA Lysis Buffer (Zymo Research) supplemented with 0.5% Antifoam DX, executed in two consecutive 5-min cycles to optimize lysate recovery. The homogenate was cleared at 12,000×g, and the resulting supernatant underwent a sequential organic extraction using phenol-chloroform-isoamyl alcohol (25:24:1) followed by chloroform-isoamyl alcohol (24:1), with vigorous 15-min agitation periods. The final aqueous phase was isolated, mixed with an equal volume of absolute ethanol, and purified using Zymo-Spin IIICG columns (Total RNA Purification Kit, Zymo Research). To eliminate genomic DNA contamination, an on-column DNase I treatment was performed for 30 min at room temperature. High-quality total RNA was subsequently eluted in 40 μL of nuclease-free water.

For analysis of RNA and protein in mitochondria, the Minute™ Mitochondria Isolation Kit for Muscle Tissues/Cultured Muscle Cells was used following the manufacturer’s protocol.

#### Western blot analysis

Total protein lysates were prepared in RIPA buffer containing protease inhibitors. Proteins were size-separated by SDS-PAGE and transferred onto nitrocellulose membrane (Life Technologies). For western blot analysis, primary antibodies were employed that recognized MYH/MHC (B-5) and HSP90 (F-8) (Santa Cruz Biotechnology), MEF2C (D80C1) (Cell Signaling Technology), Total OXPHOS Human WB Antibody Cocktail including ATP5A, UQCRC2, SDHB, COX2, NDUFB8 (ab110411) (Abcam), LRPPRC (F-7) (Santa Cruz Biotechnology), SLIRP antibody (ab51523) (Abcam), and HADHB Antibody (E-1) (Santa Cruz Biotechnology). Details in the Resources Table. After incubation with the appropriate secondary antibodies, protein signals were developed using chemiluminescence.

#### Immunocytochemistry and Confocal Imaging

Cells were fixed in a solution of 4% paraformaldehyde in 0.1 M PBS (pH 7.4) for 20 min, washed with PBS, and incubated for 5 min in a solution of 0.2% Triton X-100 and 5% normal goat serum in 0.1 M PBS containing 5% normal goat serum. Fixed cells were then incubated in the same solution containing primary antibody at 4°C for 20 h, washed with PBS, and incubated for 2 h at 25°C in PBS containing FITC or rhodamine-conjugated secondary antibody. Primary antibodies were used that recognized MYH/MHC (B-5, Santa Cruz Biotechnology), VDAC (D73D12) (Cell Signaling Technology), and LRPPRC (F-7) (Santa Cruz Biotechnology) were employed (details in Resources Table); nuclei were visualized by staining with DAPI (4’, 6-diamidino-2-phenylindole).

#### Library preparation protocol

Total RNA was isolated from cells with the Direct-zol™ RNA MiniPrep kit (Zymo Research), which includes a digestion step using DNase I. And quality check on the Agilent 2100 Bioanalyzer (RNA 6000 nano kit (Agilent 5067-1511), total RNAs (500 ng) was subjected to rRNA depletion with the Low Input RiboMinus Eukaryote System v2 (Thermo Scientific Cat# A15027). rRNA-depleted samples were then used for cDNA synthesis with Ovation® RNA-Seq System V2 (Nugen Cat# 7102) following the manufacturer’s protocol. Briefly, the cDNA first strand was synthesized from rRNA-depleted RNAs using a unique first-strand DNA/RNA chimeric primer mix and reverse transcriptase (RT) included in the kit; double-stranded cDNAs were then synthesized. After purification with Agencort RNA CleanUp XP beads, the double-stranded cDNA products were amplified with SPIA (single-primer isothermal amplification) included in the kit. The amplified products were then purified with Qiagen QIAquick PCR Purification Kit (Qiagen Cat# 28104). The purified products were checked on Agilent 2100 Bioanalyzer with the DNA 1000 kit (Agilent Cat#5067-1504) and fragmented using a Bioruptor. The fragmented cDNAs were checked again on Agilent 2100 bioanalyzer with the DNA 1000 kit.

The fragmented cDNAs were used for library preparation using the Illumina TruSeq ChIP Library Preparation Kit (Illumina Cat# IP-202-1012) following the manufacturer’s protocol. Briefly, the cDNAs were subjected to end repair, 3’-end adenylation, adapter ligation, and purification with AMPure beads (Beckman, Cat#A63881). The purified cDNAs were amplified by PCR and purified again with AMPure beads to generate final libraries. RNAseq libraries were sequenced on an Illumina Hi-Seq 2500 instrument with single-end (141 bp) at a depth of 68-80 million reads.

#### RNA-seq analysis

Single-end and double-end sequencing were performed on Illumina HiSeq 2500. The FASTQ files were extracted using bcl2fastq v2.20.0.422, trimmed for adapter sequences using Cutadapt v1.18, and aligned to human genome hg19 Ensembl v82 using STAR software v2.4.0j. FeatureCounts was used to create gene counts from six samples. The DESeq2 package version 1.26.0^70^ in R (version 3.6.3) was used to carry out the differential expression analysis. Statistical testing was performed using the Wald test and those transcripts with Benjamini-Hochberg adjusted p-values < 0.05 and absolute log_2_ fold change > 1 were considered to be differentially expressed. The RNA-seq data were deposited in GSE261476 (token oterkuicznajdij) and GSE261477 (token yzqxkuykfvopbqx).

#### Modified ChIRP analysis for mass spectrometry (MS) and western blot analyses of RNA-bound proteins

Fifty 15-cm dishes of AB678 cells were used per ChIRP (chromatin isolation by RNA purification) experiment followed by protein analysis. Cell harvesting, disruption and ChIRP were performed as previously described.^23,71,72^ Cells were crosslinked in 3% formaldehyde for 30 min, quenched with 125 mM glycine for 5 min, and incubated in lysis buffer (50 mM Tris-HCl [pH 7.5], 10 mM EDTA, 1% SDS, protease inhibitor, RNase inhibitor). Cell extracts were sonicated in Bioruptor® (On: 30 sec, Off: 45 sec, Intensity: strong) for 15 min and centrifuged at max speed (16000 × g). Supernatants were collected and pre-cleaned with streptavidin magnetic beads (Dynabeads MyOne Streptavidin C1, Thermo Fisher Scientific) for 45 min at 37°C. Supernatants were incubated with antisense oligonucleotides (ASOs) designed at https://www.biosearchtech.com/support/tools/design-software/chirp-probe-designer directed at *LANCL1-AS1* or lacZ (negative control) (Table S4) in hybridization buffer (50 mM Tris-HCl [pH 7.5], 1 mM EDTA, 1% SDS, 750 mM NaCl, 15% formamide, protease inhibitor, RNase inhibitor) for 16 h at 37 °C. Finally, the probes were pulled using streptavidin magnetic beads and then washed five times in wash buffer (2× SSC, 0.5% SDS) at 37 °C. The bound proteins were eluted and reverse-crosslinked by boiling in SDS sample buffer (95 °C for 30 min) for mass spectrometry (Table S1) and western blot analysis.

After ChIRP assay, mass spectrometry analysis was performed by Poochon Scientific. Briefly, after in-gel trypsin digestion, peptides were analyzed by liquid chromatography-coupled tandem MS (LC-MS/MS) using a QExactive hybrid quadrupole orbitrap mass spectrometer (Thermo Scientific) with a Dionex UltiMate 3000 RSLCnano system. Peptide identification and protein assembly were performed on a Thermo Proteome Discoverer 1.4.1 platform. Tandem mass spectrometry datasets were analyzed against corresponding UniProtKB/Swiss-Prot database using the SEQUEST and Percolator algorithms. The proteomics results are shown in (Table S1).

#### smFISH

Myoblasts and myotubes were grown on glass coverslips coated with poly-d lysine and then fixed, permeabilized, and washed to prepare for hybridization, as previously described.^73^ To label mitochondria, myotubes were incubated with 25 nM of MitoTracker Deep Red (ThermoFisher Scientific, M22426), prepared following the manufacturer’s instructions, for 15 min before continuing fixation. Briefly, a set of probes, each 20 nt in length and designed to bind the entire length of *LANCL1-AS1*, were ordered from LGC Biosearch Technologies with a 3’ NH2 modification. The probes were pooled, coupled with Texas Red dye, and purified by high-performance liquid chromatography (HPLC). The sequence of probes used is provided in Table S5. The coverslips were incubated with the probes diluted in hybridization buffer. After overnight incubation, the coverslips were washed and processed for immunofluorescence (IFA) using antibodies recognizing LRPPRC and TOM20 following the previously described protocol.^74^ Finally, the coverslips were stained with DAPI, and mounted on clean glass slides and the edges of coverslips were sealed with nail polish. A 100× oil objective on a Nikon TiE inverted microscope with a PIXIS:1024B CCD camera (Princeton Instruments Inc.) and Metamorph imaging software was used to acquire z-stack images. The quantification of *LANCL1-AS1* and the colocalization between *LANCL1-AS1* and mitochondrial proteins were determined using a custom-designed algorithms in MATLAB software (MathWorks Inc). The z-stacks were combined, and overlays were obtained using NIH ImageJ software.

#### Mitochondrial poly(A) tail (MPAT) assays

Mitochondrial poly(A) tail (MPAT) assay was performed using the Poly(A) Tail-Length Assay Kit (Invitrogen) following the manufacturer’s instructions. For the analysis of poly (A) tail by Gel Electrophoresis, the PCR products were further analyzed using E-Gel precast agarose gels (Invitrogen). For the analysis of poly (A) tail by Sanger sequencing, the PCR products were further subcloned using TOPO-TA Cloning (Invitrogen) following the manufacturer’s instructions. The primer sequences used to detect specific human mt-mRNAs for Poly(A) Tail-Length Assay are listed (Table S6).

#### Mitochondrial activity measurement using Seahorse

AB678 Cells were plated at a density of 2 × 10^5^ cells/well in a 24-well Seahorse tissue culture plate, 24, 48 or 72 h before the day of the assay. On the day of the assay, the culture media was changed to Seahorse Assay media (Agilent #103335-100) supplemented with 10 mM glucose, 1.0 mM sodium pyruvate and 2 mM glutamine. The media had been pre-warmed and the pH adjusted to 7.4. Cells were washed twice with assay media and assay media was added to a final volume of 525 μl per well. The plate was incubated at 37°C (no CO_2_) for 1 h prior to assay. Meanwhile, a hydrated, 24-well flux plate was prepared using the XF Cell Mito Stress Kit (Agilent #103015-100); 75 μl of 8 μM Oligomycin was added to port A, 75 μl of 45 μM FCCP was added to port B, and 75 μl of 5 μM Rotenone/Antimycin A was added to port C for final concentrations of 1.0 μM, 5.0 μM and 0.5 μM, respectively. The XFe24 Flux analyzer was programmed to take 3 basal measurements, 3 min apart, followed by injection of Port A, with 2 cycles of mixing for 3 min, 2 minutes waiting, and measurement for 3 min. This cycle was repeated for Port B and Port C. The results were analyzed using the Agilent Wave 2.6.1 software.

### QUANTIFICATION AND STATISTICAL ANALYSIS

The results are represented as the means ± SEM. Statistical comparisons of the results were evaluated using Student’s t-test. Statistical significance was indicated as follows: *, p<0.05; **, p< 0.01); and ***, p< 0.001.

