## Supplementary Text and Figures for "Reduced *LANCL1-AS1* in old human skeletal muscle diminishes mitochondrial activity, shortens mt-mRNA poly(A) tails, and suppresses myogenesis"

### SUPPLEMENTAL INFORMATION

#### LEGENDS FOR SUPPLEMENTAL FIGURES

**Supplemental Figure 1. Annotation of skeletal muscle lncRNA *LANCL1-AS1*, related to Figure 1.**

**(A)** The levels of *LINC00667* (and loading control *GAPDH* mRNA for normalization) were quantified by RT-qPCR analysis in RNA collected from myoblasts undergoing myogenesis, collected at the times shown.

**(B)** Levels of *LANCL1-AS1*, as quantified by RT-qPCR analysis from RNA prepared from human cervical carcinoma HeLa cells, WI-38 human diploid fibroblasts, mouse embryonic fibroblasts (MEF), human vascular smooth muscle cells (VSMC), human monocytic leukemia (THP-1) cells, human proliferating (Prolif.) and differentiated (Diff.) AB678 myoblasts, and human proliferating and differentiated SH-SY5Y neuroblastoma cells; data were normalized to the levels of *Gapdh* mRNA (MEF) or *GAPDH* mRNA (all other cell types). Expression levels of the muscle-specific transcript *MYL1* mRNA in the same cell types.

**(C)** Schematic depicting the difference between *LANCL1-AS1* expressed in AB678 skeletal myoblasts and the NCBI reference sequence for *LANCL1-AS1* (NR_110604.1).

**(D)** Sequence of skeletal muscle *LANCL1-AS1* as determined by 5' RACE analysis of differentiated AB678 myoblasts, depicting a novel 175 nt insertion (red) detected in AB678 myoblasts.

**(E)** Calculation of copy number in proliferating and differentiated AB678 cultures by three different methods: standard curve assay, droplet digital PCR analysis (with ddPCR analysis image below), and equivalent Ct value by RT-qPCR (the latter comparing to *GAPDH* mRNA, which is present in ~1300 copies per cell and per nucleus equivalent in AB678).[22,29]

**Supplemental Figure 2. Silencing *LANCL1-AS1* attenuates human myogenesis, extended data, related to Figure 2.**

**(A)** AB678 myoblasts were transfected with Ctrl siRNA or LANCL1-AS1-directed siRNA #2; 24 h later, they were placed in differentiation medium for 72 h, whereupon differentiation was monitored by evaluating myotube formation using phase-contrast microscopy.

**(B)** AB1167 myoblasts were transfected with Ctrl siRNA or LANCL1-AS1-directed siRNA #1; 24 h later, they were placed in differentiation medium for 72 h, whereupon differentiation was monitored by assessing MYH levels (green) by immunofluorescence.

**(C)** AB1167 myoblasts were transfected with Ctrl siRNA or LANCL1-AS1 siRNA #1. The fusion index and number of nuclei (stained using DAPI) per myotube after differentiation for 72 h were quantified.

**(D)** AB1167 myoblasts were transfected with Ctrl siRNA or LANCL1-AS1 siRNA #1. The levels of *LANCL1-AS1* 48 h later were calculated by RT-qPCR analysis.

**Supplemental Figure 3. Extended RNA-seq analysis after silencing *LANCL1-AS1*, and mitochondrial activity during myogenesis, related to Figure 3.**

**(A)** Heatmap representation of the changes in the transcriptomes associated with biological processes after silencing *LANCL1-AS1* (complements data in Figure 3C). RNAs encoded by mitochondrial genes (from MitoCarta3.0 dataset) that changed >2-fold (padj<0.05) 48 h after silencing *LANCL1-AS1* were included.

**(B)** Schematic of genes on the mitochondrial genome, including those from which mt-mRNAs are transcribed. Created by modifying a template provided by BioRender.

**(C)** Mitochondrial activity measured before the start of myogenesis (time 0 h), at 24 h into myogenesis, and at 72 h of differentiation. Green rectangle, maximal respiration; ATP production, pink rectangle. OCR was measured using the Seahorse XFe24 analyzer. Data in (C) are the means ±SEM of three independent replicates.

**Supplemental Figure 4. Interaction of *LANCL1-AS1* with LRPPRC, related to Figure 4.**

**(A)** To complement the ChIRP-MS experiment (Figure 4A,B), RT-qPCR analysis was performed to monitor the enrichment of *LANCL1-AS1* in each biotinylated ASO pulldown group.

**(B)** To complement the ChIRP-western experiment (Figure 4E), the ability of the ASOs to pull down *LANCL1-AS1* was evaluated by RT-qPCR analysis. Fragment I was detected by primers 1-4, fragment II by primers 5-8, fragment III by primers 9-12, and fragment IV by primers 11-14. 'All' was detected by RT-qPCR analysis using primers similar to those used to detect *LANCL1-AS1* in Figure S2D**.**

**(C)** Quantification of smFISH puncta per myoblast (2.3 ± 0.48 copies per cell) and myotube (43.4 ± 6.4 copies per nucleus equivalent).

**(D)** Representative smFISH analysis of *LANCL1-AS1* colocalizing with MitoTracker, LRPPRC, TOM20. *Far left*, DIC micrographs. *Left center*, RNA signals using probes for *LANCL1-AS1* RNA using a gray scale to represent RNA; empty red circles highlight the specific RNA signals. *Right center,* RNA signals using probes for *LANCL1-AS1* RNA merged with MitoTracker signals using gray scale to represent RNA and yellow arrows to identify the co-localization of *LANCL1-AS1* and MitoTracker. *Far right*, RNA signals using probes for *LANCL1-AS1*, merged with LRPPRC or TOM20 signals using antibodies recognized LRPPRC or TOM20, using gray scale to represent RNA and yellow arrows to identify the co-localization of *LANCL1-AS1* and mitochondrial proteins. Scale bar, 5 μm.

**(E)** Representative smFISH analysis for *LANCL1-AS1* colocalizing with LRPPRC; *left*, DIC micrographs; *center*, merged RNA signals (red) using probes for *LANCL1-AS1* RNA (*top*) or the control nuclear lncRNA *lncFAM* (*bottom*), merged with protein signals using antibodies against LRPPRC (green) and DAPI (blue).

*Right*, orange spots denote where lncRNA signals colocalize with LRPPRC protein, as identified using MATLAB. Scale bar, 5 µm.

**(F)** AB678 myoblasts were transfected with Ctrl siRNA or siRNA directed at PNPase (*PNPT1* mRNA); 24 h later, they were placed in differentiation medium for 72 h, whereupon differentiation was monitored by evaluating myotube formation using phase-contrast microscopy.

**Supplemental Figure 5. Silencing *LANCL1-AS1* reduced the poly(A) tail length of mt-mRNAs, extended data, related to Figures 5 and 6.**

**(A)** AB678 myoblasts were transfected with Ctrl or LANCL1-AS1 siRNAs; 24 h later, they were placed in differentiation medium for an additional 24 or 48 h, whereupon total DNA was extracted and the levels of mt-DNAs at 24 and 48 h after inducing differentiation were quantified by qPCR analysis and represented using a heatmap. *Right*, specific measurements of the levels of *mt-ND1* DNA, as assessed by qPCR analysis, are shown.

**(B)** By 24 h after transfecting Ctrl or LANCL1-AS1 siRNAs, AB678 myoblasts were differentiated for an additional 72 h, whereupon cells were collected and mt-mRNA poly(A) tail length (MPAT) assay performed. An example of Sanger sequencing performed to evaluate the length of the poly(A) tail for an mt-mRNA (*mt-ND2* mRNA) in one of the clones is shown.

**(C)** By 24 h after transfecting Ctrl or LANCL1-AS1 siRNAs, AB678 myoblasts were differentiated for an additional 72 h. Cell lysates were then subjected to co-immunoprecipitation (co-IP) assays using an antibody that recognizes LRPPRC or a control IgG. The levels of SLIRP and LRPPRC in the IP materials were then evaluated by western blot analysis; signals in 'Input' lysates, without IP, were also assessed.

**(D)** Complete set of MPAT assays (as described in Figure 6B) conducted to determine the poly(A) tail length of mt-mRNAs from muscle biopsies taken from young and old human skeletal muscle. MPAT samples were analyzed by monitoring mobility shifts of DNA fragments during electrophoresis on agarose gels.

**(E)** Search for sequences most similar to human *LANCL1-AS1* in other species using NCBI BLAST. Orange arrow points to *Macaca mulatta*, bearing 79% homology (*top*). Schematic depicts BLAST comparison of human and monkey *LANCL1-AS1* (*bottom*).

#### LEGENDS FOR SUPPLEMENTAL TABLES

**Supplemental Table S1.** LANCL1-AS1 Annotation of skeletal muscle lncRNA *LANCL1-AS1*.

**Supplemental Table S2.** siRNAs used in this study to silence different RNAs.

**Supplemental Table S3.** Oligomers used in this for amplification in RT-qPCR analyses.

**Supplemental Table S4.** ASOs used in ChIRP analysis.

**Supplemental Table S5.** Oligonucleotide probes used in smFISH.

**Supplemental Table S6.** Primers used to measure the length of poly(A) tails in mt-mRNAs (MPAT assays).

A

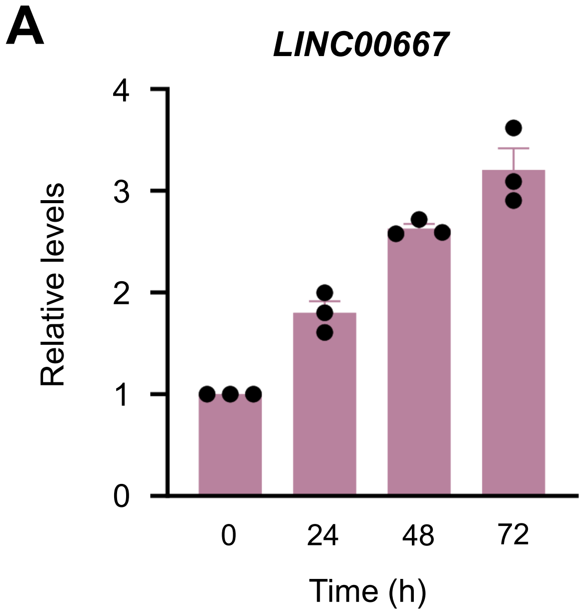


B

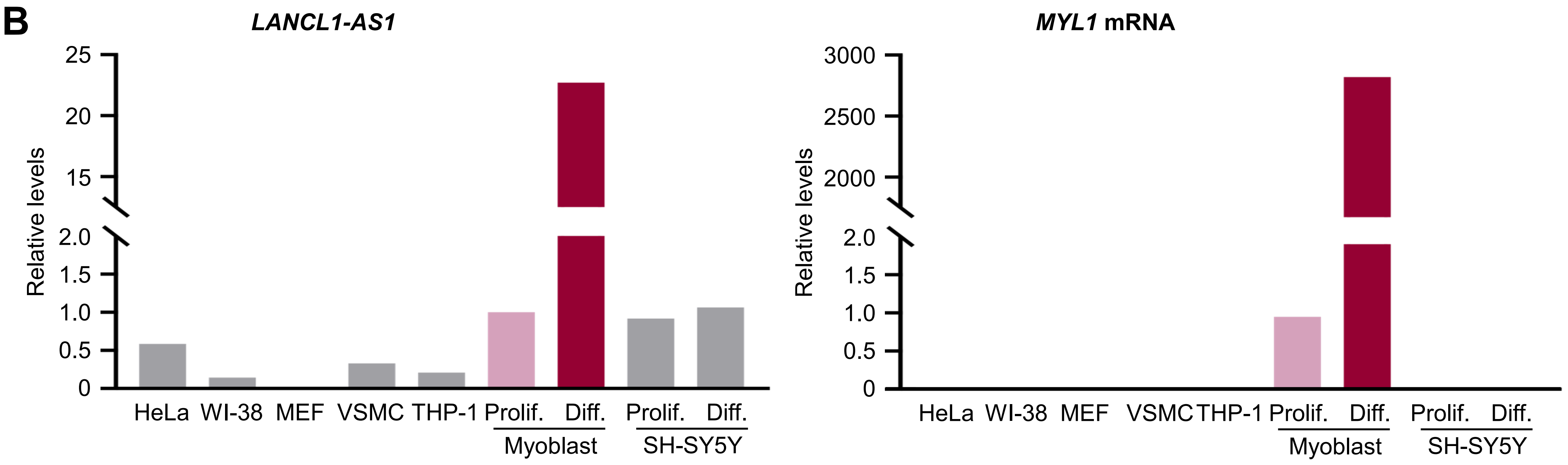


C

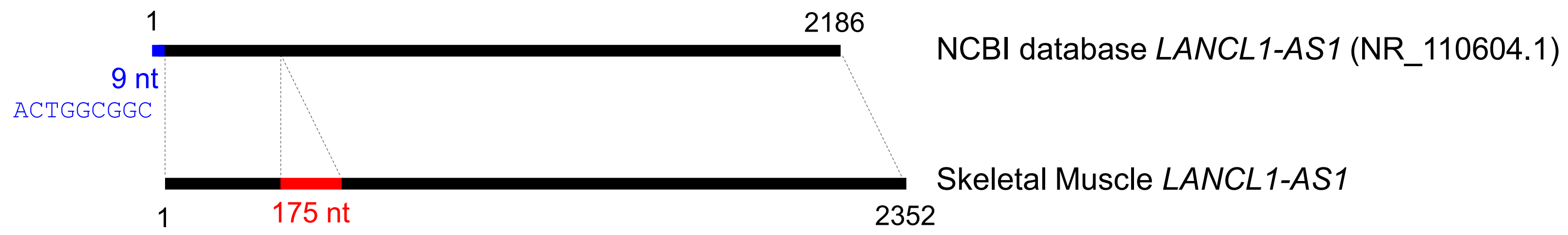


D

CTCAGTCCTTGTGTTTGGAGCAGTTCTGCTAGAGGAGGAAACAGCACAAAGGGCACTCTTATGCTGCTGTAAACCAGTGCCAGGATTTTTAAAATGTGAGACACCTGTTACCCTGTGCCCAGATT
TCATATTGTTCAAACACACATGCTCTACATACAATTTGTGCAGTTAATGCAATCATCACAAGGTCCTAGGCCGACCTACAGCCTCAGCTTACGAAGATGACAAGATTAAGAGATTAAAGTAAAGA
CAGGCATAAGAAATCAAAAGAGTATTGATTAAGGAAGTGAGAGTGTGAAGAGGAGGAGAAAGGCTGAAGAGGAGGAGAAAGGCTAAGATCTTTTCCAATGCAAAATTAGGAAAAAAATATACTACA
GGTAAGATGGAGCAAGCATGCTCTGCACCTTGTGTTCCCTCAAAATGTGGCTCTAAAACCTGGACAGGATACAGCAGCTATTTGAGGTCTCTGAAAAGCAAACAGTAACATGCAAAAAGGGGAAG
AGGACAGGAATCTGAAGTAGCACCAAACCAATGATTTATGAAAAAATAAGAAGTGAAAGTATTCTCTGACTGCAAGTATGACCACCTGTCTTTCTATATCAGAACCCAAGCTAAAAGACTGCATT
TACTACCCTTCCTTGCTAACAAGAGTGACTACGTGACTATGTTCTGGCCAATGCAAACAGCCTGAAAAGCTGATGTAATGACTGGATCTTCAGCAGCCACCTTGAGCAAGAGGTAGAAGCTAGAG
GCTGAGGAATGATGGAGCAGAAAGGAAGGAGCTGGGGTTCCTATTAACATCCTGAAGCTACACTATCAGGCCTGGACTGACCAACTCTTTCTGGCCTTCTTTTACAAGAGAAAAATACAGTTATT
TAAACCACTATACTTCTGTTTTCTTTTTTGGGGTATAGATAACTGAACCTAATCCTGTCCCAAATCCTGATCCTATCTAACCAGTGTTAGCTATCAGCCCATACGTATACTGGAGAATGACGTTT
AACAAAATAAGATGTTAAAGTTCTAACTAATTGTAATAATCATGCAGTTTTCTCATGCTGAACAAAGTTAAAGAAATATAATGTGTGTTTCCTAAAGAGTGTCATTTTTTGGAAAGATAACATGG
GGCACAAAGTAACATAAAAATCTACATATAATTATGTAAACATTGCTTTGACATACAACACCAAATATTTTTATTGTATTTGTGAATGCATTTCTTGGTAATAAAACGGCAGTTATGAATAAGTG
ATACTTGGAAAGCTATAACATAATGGGAGGAGAAGGTATTAACACTGTGATTTTATGGGGTTGAATCAGTTTTAGTTTCTTATACAACTCAGAGCTTTACGTTATTTAGACTGTCTAAACACTGG
CGCAAGGTAGCATTTTACTTATCAAAACACTGGAGGAGGCAGTCTATTTTATGTGGGAATAATAGCTTTTTAACAGTGTTCGATTAGGAGACATTAGGAACCTTGCCTTGAAAACCATTAGCAAA
TTTATCTAGAAGATTTTACGTTACATCTTAAAACAACCAGCTGCTAAGGACATCCTAAACTGCTTAGTATGCCAATTTGCCTCAGATGCTTGCCATATAAGCTGCTTTCAGGAGTCAGCAATGAG
TATCATAGATACTTGAGGATAAAAGGTTGAAGGATATTGGCATTCAAAGATGAAGCCAGAAACCCTGTACAGCCAAAAAAATTAAACTTGCATTTTACTTTGGAGTGAAATGTCTTTGCTATTTC
CAAAGTTTTTAACTTTTGGGTTCTCAACTAGCCCTAGCAATATAAGGGCCTGACTAATGACTCTTCCCAACATTGTCAAACTTAATTCAAACTTGAAATCAATCCCCTTTTTTTTCCTCCTAAAG
CAAAGTAACTCACAATAATAATATGAGTACATGTGAATGGCCAATTTAGAAGTTTAAGTCATTCTTTTGTAATCAGTACTAGAATTAACTAGTGATTTTTTTTTTTGGCAAGGAATAACAAAATA
ATAGAAACTCCATATATATTAGAAAAAGATCTGTAAGTTTGAGCCTTATATATTATTACAATGTGGCAATCAAGAGGAAAACTCAAGACAACTGTGCGCTGTGCTTGTCCTGTTTTGTTCAAACC
ACGATAATTTTAAATAAATAGTAATGTTCAACATTCACTGATGACATTATCTATCAACCACATCACACCTGGAACACAGTAGATAATAAATGGCAGCTGTTATACCTTCCTTCTTGAAACAAAAT
CCTGTCATTTCATTCTCATTTTAACTTTGTGTTAAGCTTCTACAATCCATAAAAATGATAAAACAATAAAATGGTCCAACATTAAAAAAAAAAAAAAAAAAAA

E

**Standard Curve**

| Time | *LANCL1-AS1* copy number |
|---|---|
| 0 h | **1.1** |
| 24 h | **17.8** |
| 72 h | **115.1** |

**ddPCR**

| Time | *LANCL1-AS1* copy number |
|---|---|
| 0 h | **1.7** |
| 24 h | **11.1** |
| 72 h | **84.8** |

**RT-qPCR**

| Time | *LANCL1-AS1* copy number |
|---|---|
| 0 h | **1.8** |
| 24 h | **10.6** |
| 72 h | **74.8** |

Considering *GAPDH* mRNA abundance as being 1300 copies per cell and per nucleus equivalent

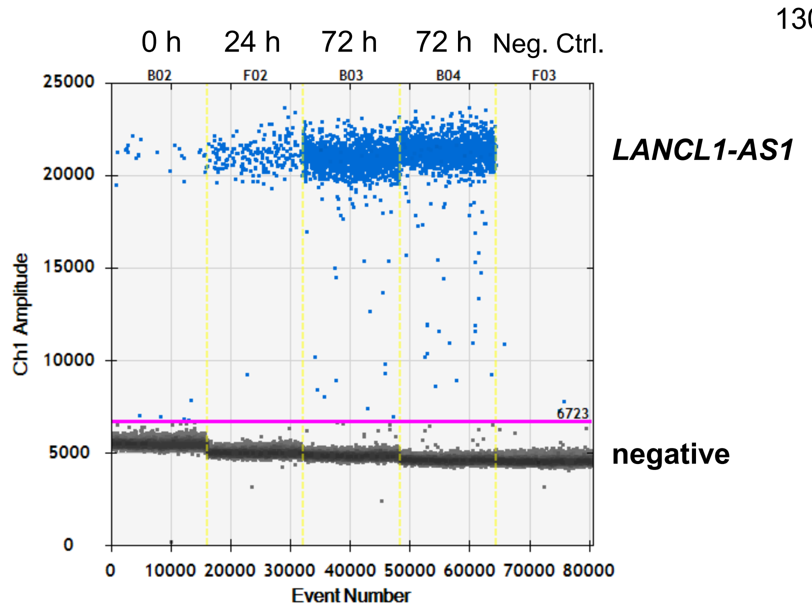

Yang et al. Figure S2

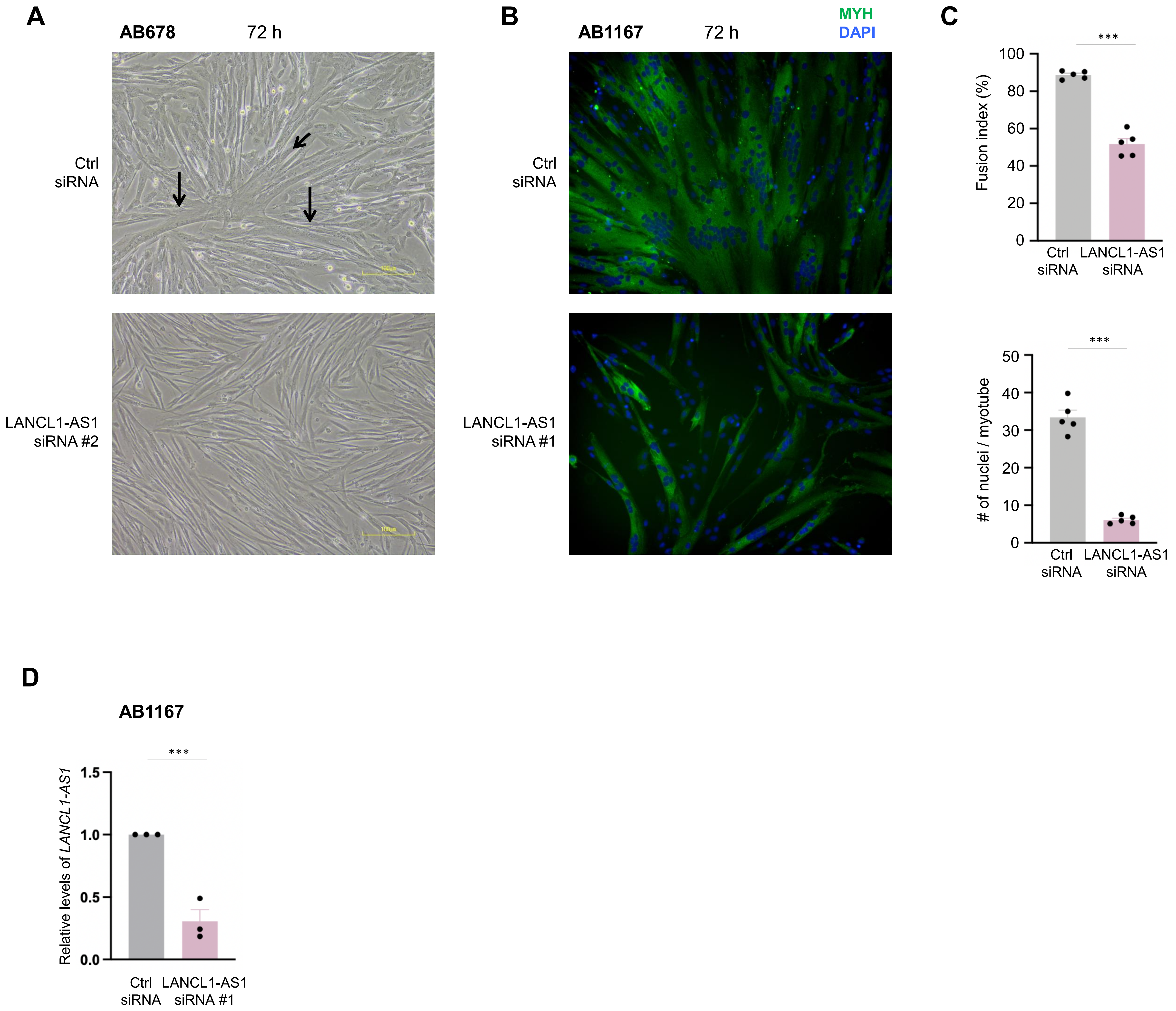

A

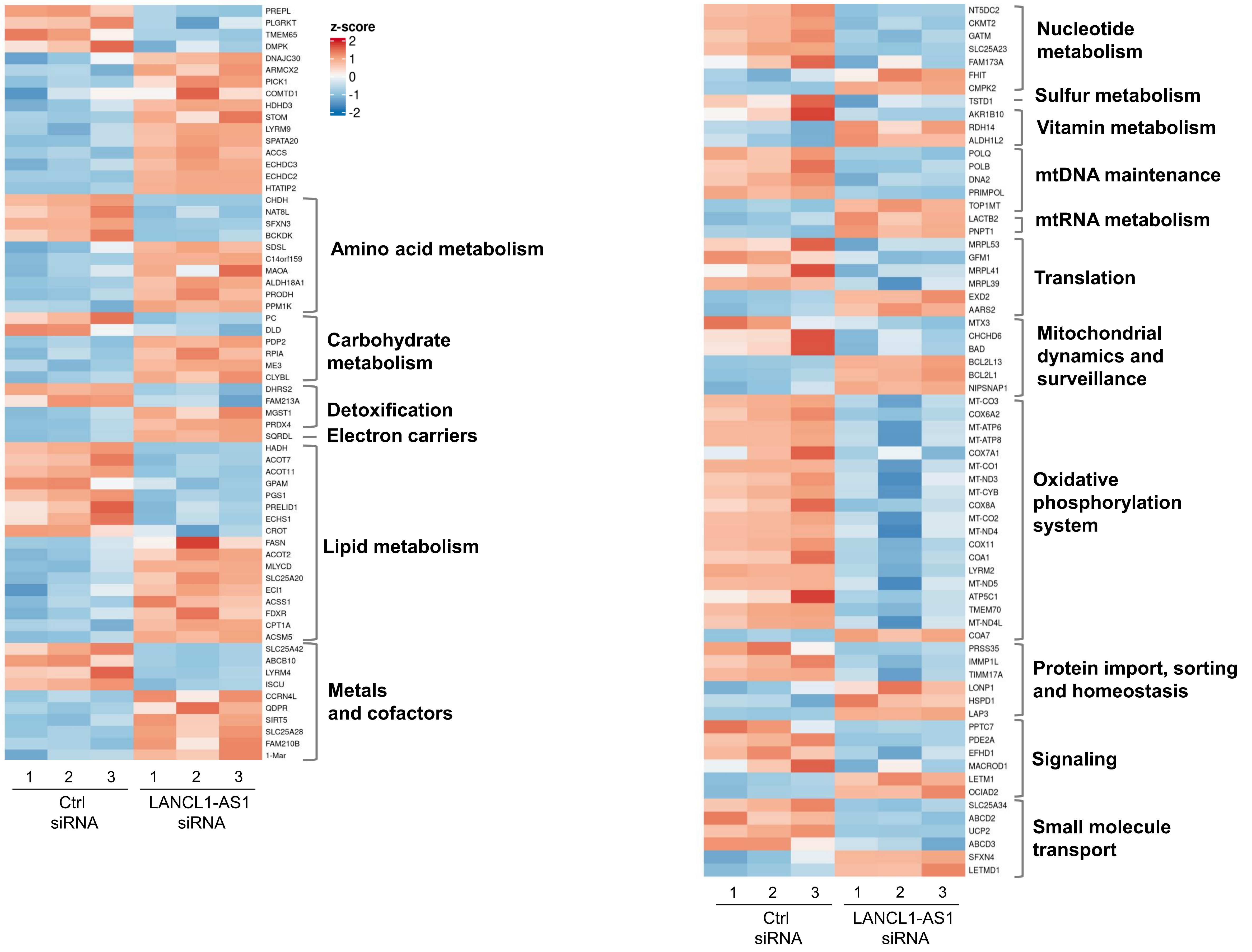
z-score
2
1
0
-1
-2
PREPL
PLGRKT
TMEM65
DMPK
DNAJC30
ARMCX2
PICK1
COMTD1
HDHD3
STOM
LYRM9
SPATA20
ACCS
ECHDC3
ECHDC2
HTATIP2
CHDH
NAT8L
SFXN3
BCKDK
SDSL
C14orf159
MAOA
ALDH18A1
PRODH
PPM1K
Amino acid metabolism
PC
DLD
PDP2
RPIA
ME3
CLYBL
Carbohydrate metabolism
DHRS2
FAM213A
MGST1
PRDX4
Detoxification
SQRDL
Electron carriers
HADH
ACOT7
ACOT11
GPAM
PGS1
PRELID1
ECHS1
CROT
FASN
ACOT2
MLYCD
SLC25A20
ECI1
ACSS1
FDXR
CPT1A
ACSM5
Lipid metabolism
SLC25A42
ABCB10
LYRM4
ISCU
CCRN4L
QDPR
SIRT5
SLC25A28
FAM210B
1-Mar
Metals and cofactors
1 2 3 1 2 3
Ctrl siRNA
LANCL1-AS1 siRNA
NT5DC2
CKMT2
GATM
SLC25A23
FAM173A
FHIT
CMPK2
Nucleotide metabolism
TSTD1
Sulfur metabolism
AKR1B10
RDH14
ALDH1L2
Vitamin metabolism
POLQ
POLB
DNA2
PRIMPOL
TOP1MT
mtDNA maintenance
LACTB2
PNPT1
mtRNA metabolism
MRPL53
GFM1
MRPL41
MRPL39
EXD2
AARS2
Translation
MTX3
CHCHD6
BAD
BCL2L13
BCL2L1
NIPSNAP1
Mitochondrial dynamics and surveillance
MT-CO3
COX6A2
MT-ATP6
MT-ATP8
COX7A1
MT-CO1
MT-ND3
MT-CYB
COX8A
MT-CO2
MT-ND4
COX11
COA1
LYRM2
MT-ND5
ATP5C1
TMEM70
MT-ND4L
COA7
Oxidative phosphorylation system
PRSS35
IMMP1L
TIMM17A
LONP1
HSPD1
LAP3
Protein import, sorting and homeostasis
PPTC7
PDE2A
EFHD1
MACROD1
LETM1
OCIAD2
Signaling
SLC25A34
ABCD2
UCP2
ABCD3
SFXN4
LETMD1
Small molecule transport
1 2 3 1 2 3
Ctrl siRNA
LANCL1-AS1 siRNA

B

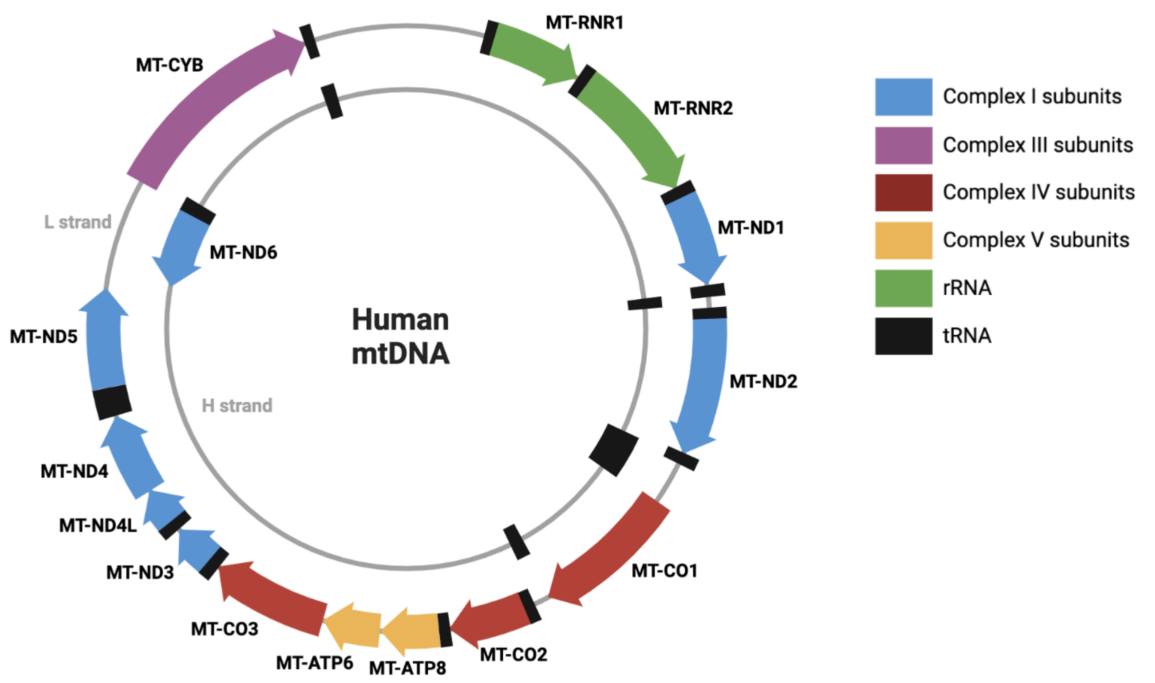
MT-RNR1
MT-RNR2
MT-ND1
MT-ND2
MT-CO1
MT-CO2
MT-ATP8
MT-ATP6
MT-CO3
MT-ND3
MT-ND4L
MT-ND4
MT-ND5
MT-ND6
MT-CYB
L strand
H strand
Human mtDNA
Complex I subunits
Complex III subunits
Complex IV subunits
Complex V subunits
rRNA
tRNA

C

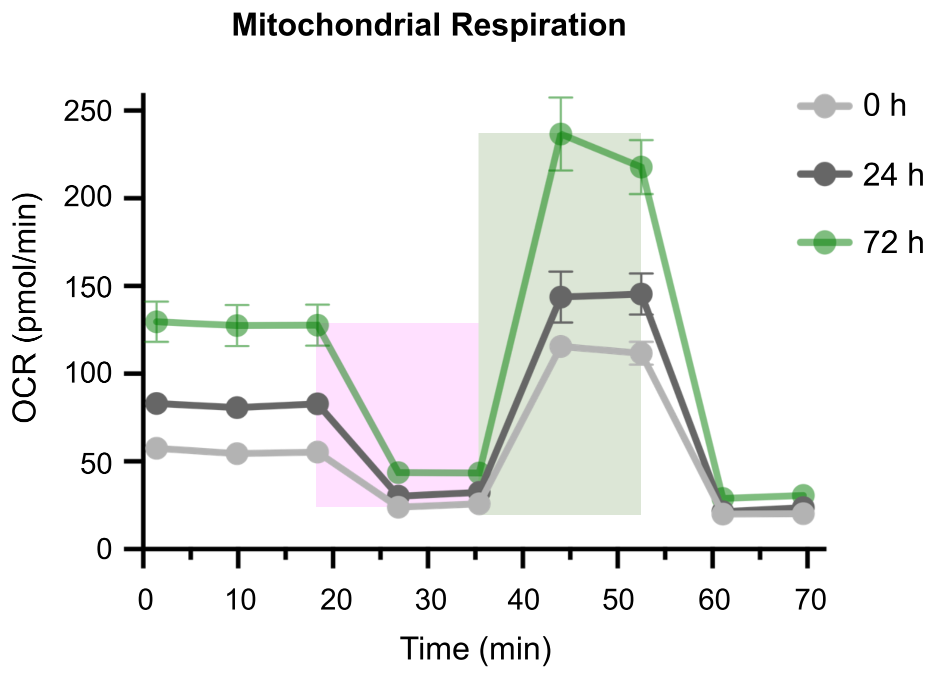
Mitochondrial Respiration
OCR (pmol/min)
0
50
100
150
200
250
0
10
20
30
40
50
60
70
Time (min)
0 h
24 h
72 h

Yang et al. Figure S4

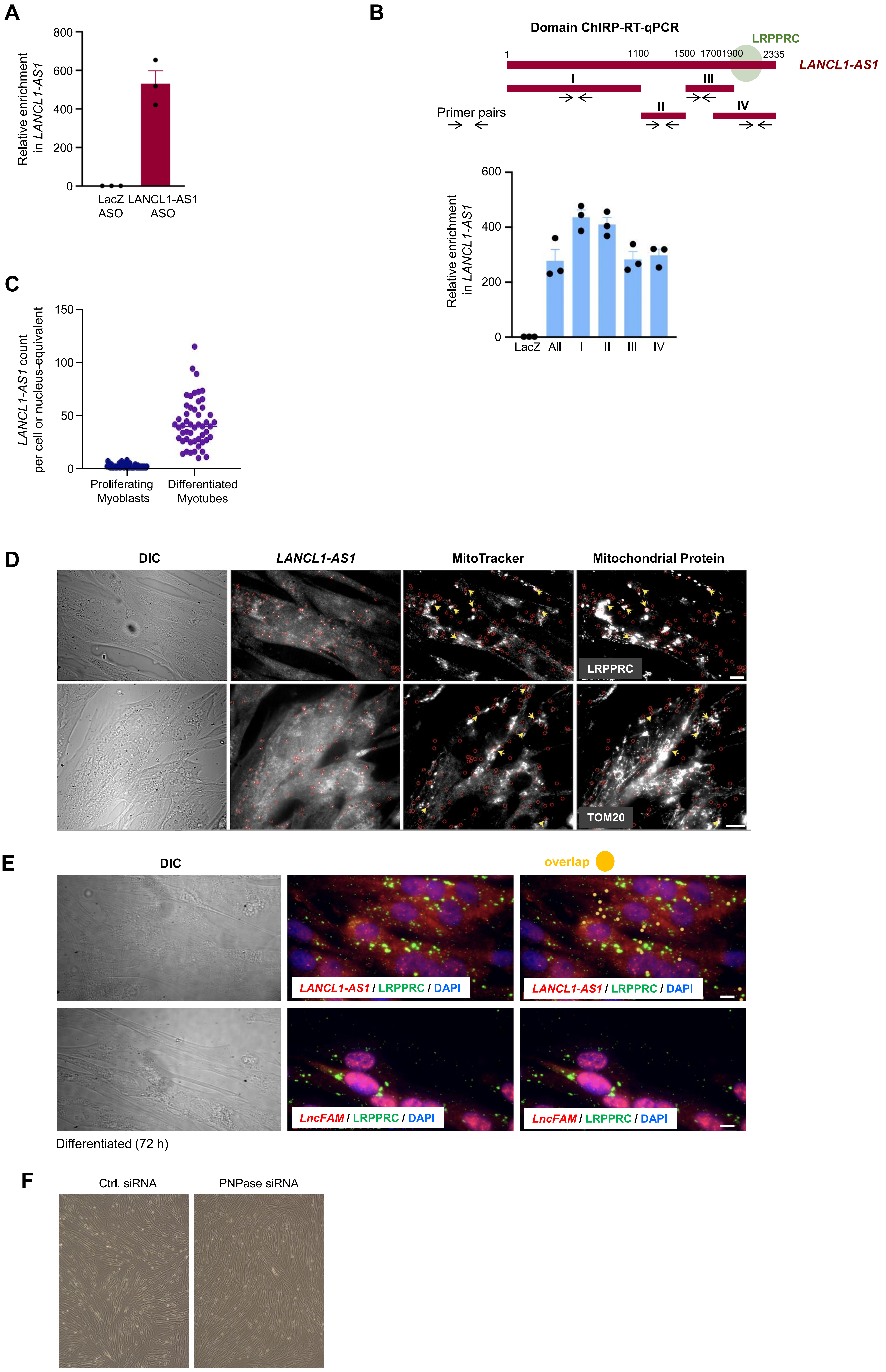

**Yang et al. Figure S5**

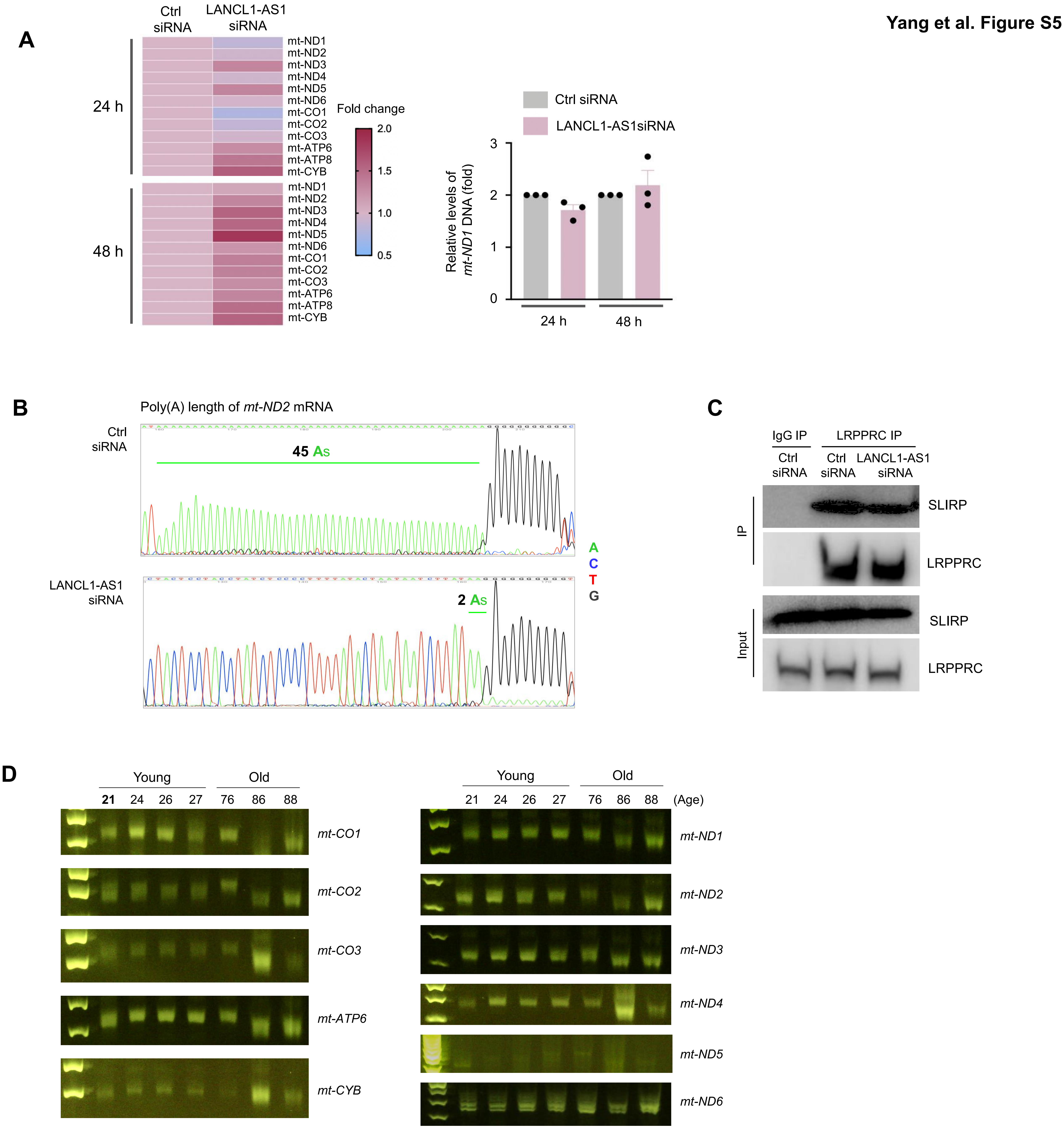


**E**

| | Description | Scientific Name | Max Score | Total Score | Query Cover | E value | Per. Ident | Acc. Len | Accession |
|---|---|---|---|---|---|---|---|---|---|
| ☐ | Homo sapiens LANCL1 antisense RNA 1 (LANCL1-AS1), transcript variant 1, long non-coding RNA | Homo sapiens | 4037 | 4037 | 100% | 0.0 | 100.00% | 2186 | NR_110604.1 |
| ☐ | Homo sapiens LANCL1 antisense RNA 1 (LANCL1-AS1), transcript variant 2, long non-coding RNA | Homo sapiens | 3862 | 3862 | 100% | 0.0 | 98.76% | 2159 | NR_110605.1 |
| ☐ | Homo sapiens LANCL1 antisense RNA 1 (LANCL1-AS1), transcript variant 3, long non-coding RNA | Homo sapiens | 3230 | 3700 | 91% | 0.0 | 99.94% | 2001 | NR_110606.1 |
| ☐ | Homo sapiens BAC clone RP11-485G2 from 2, complete sequence | Homo sapiens | 3051 | 3807 | 93% | 0.0 | 100.00% | 138890 | AC007970.3 |
| ☐ | PREDICTED: Gorilla gorilla gorilla uncharacterized LOC109025962 (LOC109025962), ncRNA | Gorilla gorilla g... | 3029 | 3029 | 79% | 0.0 | 98.21% | 1735 | XR_002004925.3 |
| ☐ | PREDICTED: Macaca mulatta uncharacterized LOC114671408 (LOC114671408), ncRNA | Macaca mulatta | 2209 | 2209 | 79% | 0.0 | 89.70% | 1839 | XR_003721402.1 |

#### Distribution of the top 1 Blast Hits on 5 subject sequences

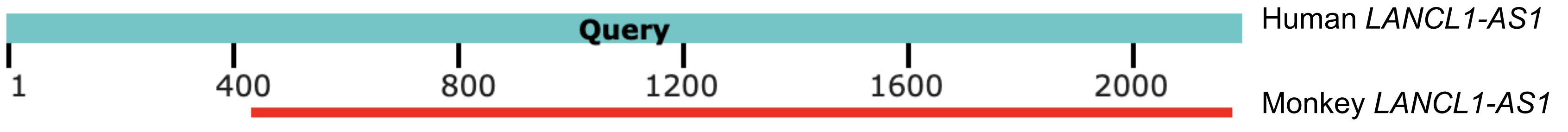
